# Phase separation determines treadmilling-like movement of actin bundles

**DOI:** 10.1101/2025.09.17.676269

**Authors:** Timon Nast-Kolb, Beatrice Nettuno, Davide Toffenetti, Moritz Striebel, Erwin Frey, Andreas R. Bausch

## Abstract

Cellular motility relies on the dynamic turnover of actin filaments, which treadmill through continuous polymerization at the barbed end and disassembly at the pointed end. Yet, the underlying physical principles remain poorly understood. A long-standing challenge has been experimentally reconstituting stable, persistent treadmilling leading to higher-order actin organization. Here, we reconstitute a minimal in vitro system, in which liquid-liquid phase-separated condensates of zyxin and VASP balance cofilin-driven disassembly to enable persistent treadmilling-like movement of actin bundles. The condensates crosslink filaments into dynamic bundles while promoting barbed-end polymerization. The localized stabilization based on condensate-mediated bundling competes against the activity of cofilin and CAP1, enabling selective disassembly at pointed ends and recycling of monomers. To elucidate the physical basis underlying this emergent behavior, we complement our experiments with agent-based simulations that quantitatively recapitulate key experimental findings. This combined approach demonstrates that an intermediate range of zyxin-VASP self-affinity, driving the phase separation, enables robust treadmilling-like movement. A weakened zyxin-VASP self-affinity fails to stabilize the bundles, whereas an excessive affinity impedes filament dynamics. Experimental reconstitution and theoretical modeling together reveal a physical mechanism by which the material properties of multivalent protein condensates govern cytoskeletal turnover, and suggest a general design principle by which biomolecular condensates can spatiotemporally organize cytoskeletal structures.

## 1 Introduction

The ability of cells to constantly reshape for movement, division, and interaction with their environment critically relies on the ability of cytoskeletal filaments to build dynamically reorganizing higher-order structures [1]. This dynamic reorganization relies on polar treadmilling, based on the underlying concept of a continuous flux of subunits by self-assembly at one end and opposing disassembly [2]. This results in filament motility that, as a core principle of active filament systems, suffices to form higher-order structures based on the local interactions of the filaments [2–5]. In eukaryotic cells, actin filaments assembling at the membrane drive motility and dynamics. Actin treadmilling was first characterized at the level of isolated filaments without additional proteins [6–8], however to ensure enhanced, persistent and localized dynamics, a complex interplay of actin-binding proteins further regulates this process [1, 9]. Despite its importance, reconstitution of actin filament treadmilling within dynamic higher-order structures has remained a central open challenge due to the complexity of the competing assembly and disassembly processes [10]. Research has mostly focused on the isolated activity of assembly or disassembly of filaments. It was found that specific barbed end polymerizing proteins like VASP (vasodilator-stimulated phosphoprotein) accelerate polymerization [11, 12], while cofilin was found to facilitate the disassembly by fragmenting and depolymerizing filaments [13–17]. The disassembly is synergistically enhanced by cyclase-associated protein 1 (CAP1) [18–20], which enables replenishment of a polymerizable monomer pool [21, 22].

While these activities are necessary for enhanced turnover dynamics, an unrestricted disassembly counteracts structural memory required for higher-order organization and additionally prohibits a purely concentration-based balance of treadmilling rates [20]. So far, this has limited reconstitution of turnover to systems with spatial separation of disassembly and continuous nucleation of assembly at solid supports without directly coupled assembly and disassembly rates [23–28]. In turn this immobilized origin of nucleation prevents filament motility and formation of higher-order dynamic actin structures. In contrast, a mechanism favoring higher-order assemblies is the direct inhibition of cofilin activity by an increase in stiffness of filaments. This is achieved in bundled filaments reinforcing formation of locally ordered filaments. However, decoration with traditional crosslinkers renders actin bundles largely resistant to cofilin activity, effectively arresting filament turnover [29–34].

An alternative mechanism for the reconstitution of dynamic higher-order bundled actin could be leveraging liquid-liquid phase separation of actin-binding proteins. In such systems, bundle stability is governed not only by weak adhesion to actin but also by the self-affinity and cohesion of the phase-separated protein condensates. VASP has recently been shown to form condensates that simultaneously polymerize and bundle filaments [35–38]. The formation of large bundles is specifically enabled by processive, aligned polymerization [37]. Multimeric interactions with zyxin could further allow tuning the material properties of the condensates [39–41].

Here, we show that the phase separation of actin-binding elongators governs the dynamic balance between filament assembly and disassembly. In a reconstituted minimal system, we demonstrate that phase-separated zyxin–VASP condensates promote localized actin polymerization and transiently bundle filaments. The size-dependent stabilization of bundles modulates cofilin-mediated disassembly, reinforcing the formation of highly organized filaments. Together with agent-based simulations, we are able to identify an optimal regime that enables robust treadmilling-like movement of actin bundles by tuning the phase separation properties. Our results reveal how the physical organization of regulatory proteins through liquid-liquid phase separation can control active filament dynamics, suggesting a general mechanism for spatial and temporal regulation in cytoskeletal systems.

## 2 Results

### 2.1 Zyxin-VASP condensates form decorated actin bundles

A central challenge in the reconstitution of actin treadmilling in vitro is the spatiotemporal coordination of filament assembly and disassembly to obtain a sustained turnover. To avoid uncontrolled disassembly by cofilin, we aim to stabilize the filaments with crosslinkers. To overcome excessive stabilization by traditional crosslinkers preventing turnover, we exploit the fluid nature of phase-separated protein condensates to generate actin bundles that are dynamically stabilized yet retain the ability to undergo controlled disassembly. Here, we utilize the ability of VASP to simultaneously form condensates, polymerize actin, and bundle filaments, providing a tunable platform for reconstituting treadmilling-like movement under defined conditions [37, 42].

To mimic its cellular localization, VASP was recruited to lipid-bound zyxin leveraging their multivalent interactions (Extended Data Fig. 1a). Zyxin in turn was attached to a supported lipid bilayer via His-tag and Ni-NTA lipids. Zyxin’s LIM domains are known to bind its N-terminus in an autoinhibitory mechanism and additionally bind to F-actin [43]. Thus, we further investigated a truncated zyxin construct without LIM domains (zyxinΔLIM) to dissect its contribution to actin network formation and effect on phase separation of VASP.

Consistent with recent observations of the formation of zyxin-VASP droplets by liquid-liquid phase sep-aration [35], the addition of VASP to homogeneously distributed full-length zyxin (zyxinFL) or zyxinΔLIM induces the formation of high-density protein condensates (Fig. 1b, Extended Data Fig. 1b). However, the zyxinΔLIM–VASP condensates form smaller and more spherical droplets than their zyxinFL–VASP counterparts (Fig. 1b, Extended Data Fig. 1b). These heterogeneous small condensates continue to mature into well-defined droplets over an hour (Extended Data Fig. 1e,f). We further observed drastically different material properties for the different constructs. While VASP triggers a reduction in fluidity in all experiments, ZyxinFL–VASP condensates exhibit markedly lower diffusivity than zyxinΔLIM-VASP, as revealed by flu-orescence recovery after photobleaching measurements (FRAP) (Fig. 1c, Extended Data Fig. 1c). Notably, even before the addition of VASP, zyxinFL forms a concentration-dependent gel-like phase likely due to intermolecular interactions on the bilayer between the LIM domains and its N-terminus (Extended Data Fig. 1c).

**Fig. 1.**
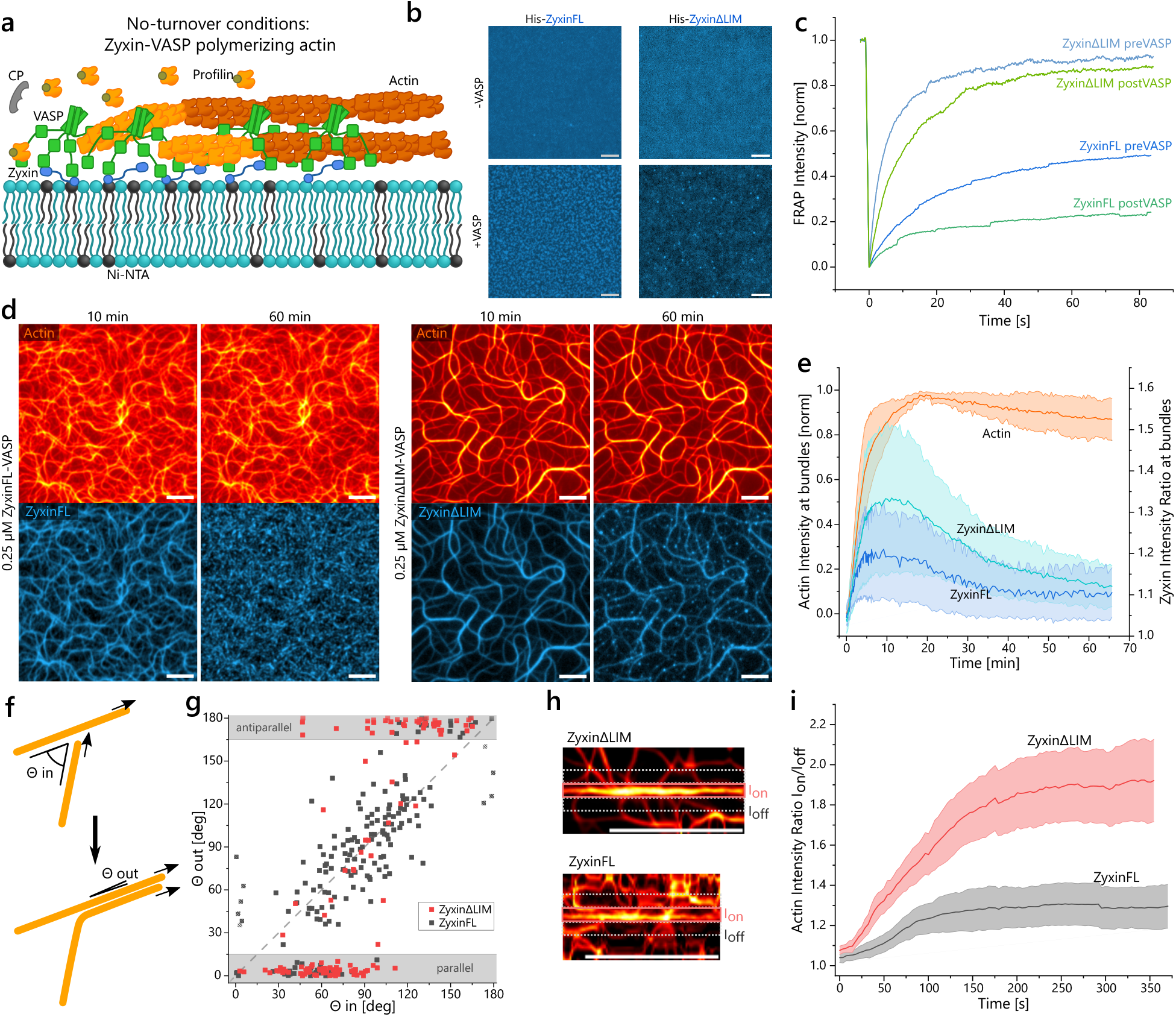
Formation of actin bundles by different zyxin-VASP condensates. **a**, Schematic illustration of the growth and decoration of actin by zyxin and VASP condensates localized to supported lipid bilayers via His-tagged zyxin. **b**, Fluorescent microscopy images of zyxin show clustering upon addition of VASP after 10 min of incubation (*C*_*zyxin*_ = *C*_*V ASP*_ = 0.25*μM* ) (scale bar, 5*μm*). **c**, Fluorescent recovery after photobleaching (FRAP) curves for the different zyxin constructs at *C* =0.25*μM* show reduced diffusivity and mobile fraction after adding VASP at *C* = 0.25*μM* . **d**, Actin TIRF microscopy images and corresponding zyxin fluorescence showing actin bundles formed by VASP bound to different zyxin constructs after *t* = 10 and 60 *min* of polymerization (scale bar, 5*μm*). **e**, Fluorescence intensity ratio on actin bundles of zyxin constructs during the polymerization (mean and standard deviation of *N* = 5 (zyxinFL) and *N* = 10 (zyxinΔLIM)). **f**, Schematic indicating the in- and out-going angles of binary interactions between actively polymerizing actin filaments. **g**, Distribution of the binary collision events. ZyxinΔLIM (*N* = 122) enables nematic alignment of filaments indicated by the grey areas showing parallel and antiparallel alignment. ZyxinFL-mediated interactions (*N* = 178) align with the dashed line, indicating no interaction. Striped squares indicate aligned filaments diverging during polymerization. **h**, Straightened actin bundles with areas indicated for on-bundle (*I*_*on*_) and off-bundle (*I*_*off*_ ) intensity (scale bar, 10*μm*). **i**, Comparison of enhanced actin polymerization intensity on the bundles between zyxinFL and zyxinΔLIM (mean and standard deviation of *N* = 5 per construct).

These differences in mobilities result in the formation of structurally distinct actin networks. Upon addition of G-actin, the condensates nucleate and polymerize filaments. Profilin and capping protein (CP) in the solution ensure that actin polymerization is specifically limited to the activity of VASP at zyxin-VASP clusters (Extended Data Fig. 2f)[9, 17, 44]. While zyxinFL-VASP polymerizes a dense and largely disordered network, zyxinΔLIM-VASP forms highly bundled actin filaments (Fig. 1d, Supplementary Movie 1). During this polymerization process, VASP’s high affinity for barbed ends [12, 37] causes the condensates to redistribute along the growing filaments, effectively decorating the network (Extended Data Fig. 2a−d). This redistribution reflects the competition between condensate surface tension mediated by its internal interactions and affinity to actin (Supplementary Note 4)[38, 45]. During polymerization, the high processivity of barbed-end binding for individual VASP proteins causes the condensates to spread along the filaments. However, once polymerization stalls out after *t* = 10 *min*, the condensate’s surface tension dominates, and the weaker affinity of VASP to F-actin-sides is insufficient to maintain a filament-bound state [12]. As a result, the system evolves toward the formation of near-spherical zyxin-VASP droplets and a progressive loss of actin decoration (Fig. 1d,e, Extended Data Fig. 2c–e, Supplementary Movie 1). Consistent with the additional internal self-interactions inferred from the FRAP data and thus a higher self-affinity, zyxinFL-VASP condensates display a faster relaxation time forming actin-unbound droplets (Fig. 1e, Extended Data Fig. 2g) and a higher actin-independent partition coefficient compared to zyxinΔLIM (Extended Data Fig. 1d). The post-polymerization decoupling of the condensates from the actin filaments further leads to a loss of bundle stabilization and filament splaying over time (Extended Data Fig. 2e).

ZyxinFL–VASP and zyxinΔLIM–VASP condensates give rise to markedly different interaction patterns between filaments polymerized by VASP on the membrane, as quantified by the angular alignment of filament encounters (Fig. 1f). In the case of zyxinFL–VASP, filaments cross without alignment. In contrast, polymerization in the presence of zyxinΔLIM–VASP promotes nematic alignment across a wide range of encounter angles (Fig. 1g, Extended Data Fig. 2h). VASP processively tracks and elongates barbed ends while simultaneously being able to bind to adjacent filaments [37]. As a polymerizing filament encounters another, continued elongation guided by the existing filament promotes local alignment and bundle formation (Extended Data Fig. 2a). This mechanism of actin bundle formation crucially depends on the mobility of VASP [37]. The low mobility of zyxinFL-VASP thus limits the processive actin elongation and nematic alignment by VASP. These alignment interactions for zyxinΔLIM–VASP on the filament level give rise to locally amplified polymerization at bundles quantified by the elevated ratio of actin polymerization within bundles (*I*_on_) to the surrounding network (*I*_off_) (Fig. 1h,i). This reinforced polymerization at bundles can not be observed for the disordered network formed by zyxinFL-VASP (Fig. 1h,i).

These results highlight how the additional interactions mediated by the LIM domains in zyxinFL–VASP condensates affect actin polymerization, filament alignment, and decoration. The difference in condensate fluidity limits both filament alignment and bundle formation. The surface tension for both constructs additionally seems to be in competition with the decoration of the actin filaments needed to maintain a bundled state. However, the weaker intermolecular interactions, and thus weaker self-affinity, in zyxinΔLIM-VASP condensates promote extended decoration of filaments and reinforced polymerization of actin bundles (Fig. 1h,i).

### 2.2 Treadmilling-like bundle movement emerges from the interplay of zyxin-VASP condensates with active disassembly

The weak and transient crosslinking provided by zyxinΔLIM-VASP condensates raises the possibility of bundles that are both structurally stabilized and continuously renewed by competition with cofilin. Thus, we examined the interplay between zyxinΔLIM-VASP condensate-mediated polymerization and bundling in the presence of profilin and CP with the disassembly enabled by cofilin and CAP1 in the reconstituted system (Fig. 2a).Cofilin and CAP1 together are expected to promote filament turnover by disassembling F-actin and recycling ADP-G-actin into polymerization-competent ATP-G-actin (Fig. 2a). However, in the first few minutes of the experiment, the high initial G-actin concentration leads to fast unimpeded polymerization, generating a dense bundled network (Extended Data Fig. 3a). During this initial stage, newly incorporated ATP-F-actin prevents binding of cofilin (Extended Data Fig. 4c). After approximately five minutes, as aging filaments accumulate, the onset of disassembly activity by cofilin leads to the emergence of directional filament dynamics. The bundle network starts displaying treadmilling-like behavior with polar growth at some ends and shrinkage at others (Fig. 2b−e, Extended Data Fig. 3a, Supplementary Movie 2). Treadmilling-like movement with discernible polymerizing and disassembling ends can be directly visualized via kymographs of individual bundles, which reveal distinct polymerization *v*_pol_ and disassembly velocities *v*_dis_ (Fig. 2d). At *t* = 30 min, a high bundle polymerization velocity can be quantified, driven by a hereto high G-actin concentration (*v*_pol_ ≫ *v*_dis_). Continued polymerization depletes the monomers, slowing down the velocity. By *t* = 1 h, the system reaches a dynamic steady state with *v*_pol_ = *v*_dis_, marking the onset of sustained actin bundle movement (Fig. 2f,g). In this state, the availability of recycled actin monomers is rate-limiting. With the initiation of filament disassembly, the actin network undergoes a pronounced coarsening of bundle size (Fig. 2h, Extended Data Fig. 3e). Individual filaments and small bundles, which are only weakly decorated and crosslinked by the zyxinΔLIM-VASP condensates, are prone to cofilin-mediated fragmentation (Fig. 2e). Concomitant recycling supplies a high ATP-G-actin pool. This in turn leads to the fast polymerization of the remaining large bundles observed at *t* = 30 min (Fig. 2f,g). Thus, this structure-selective disassembly of small actin bundles, combined with funneling of monomers towards the uncapped and processively polymerizing filaments by the high-density of VASP at zyxinΔLIM-VASP-decorated bundles, cooperate to accelerate a locally-reinforced barbed end growth. This also facilitates the emergence of large bands and vortices observable during later acquisition times (Fig. 2e,h, Extended Data Fig. 3a,e, Supplementary Movie 2).

**Fig. 2.**
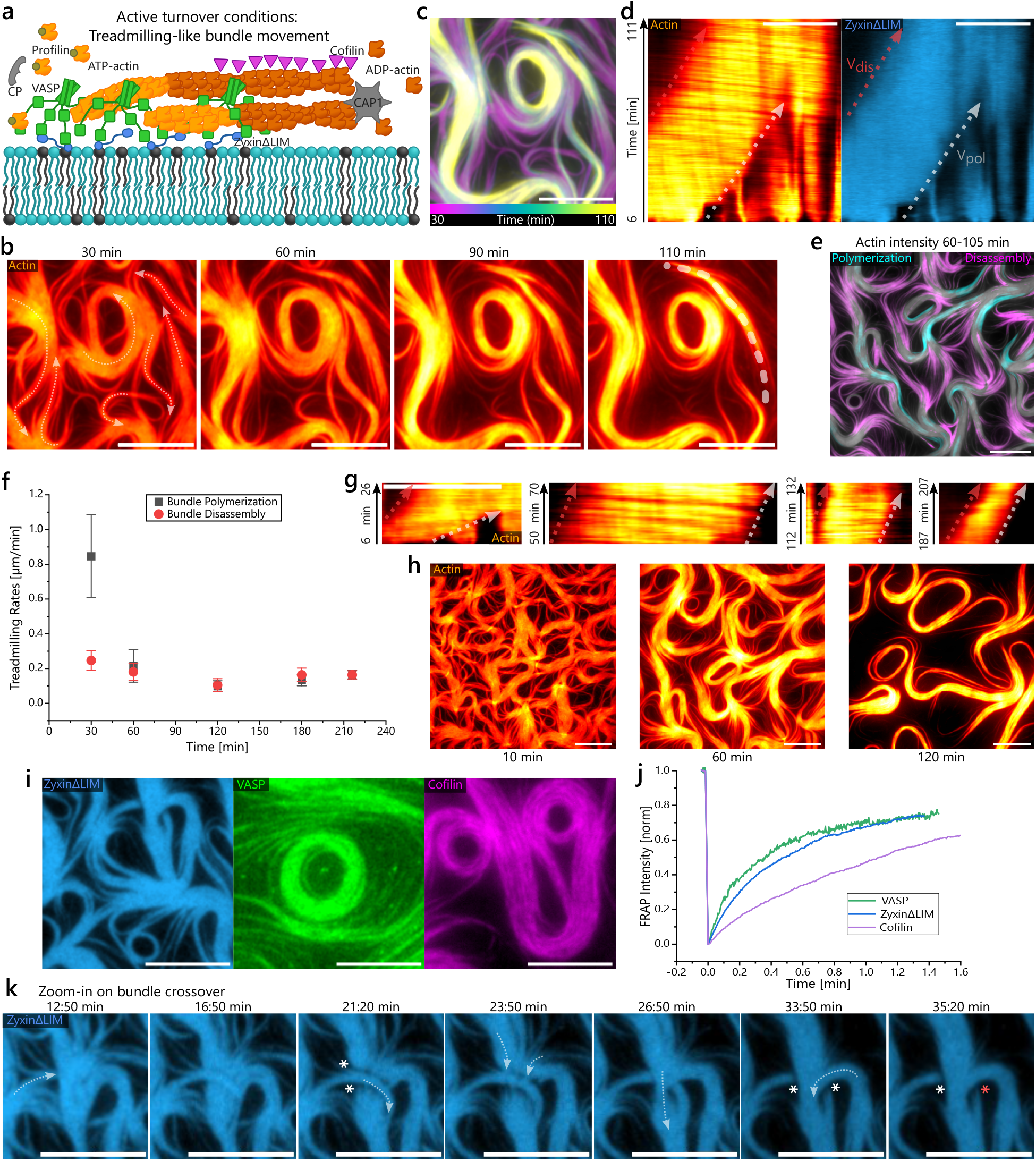
Interplay of ZyxinΔLIM-VASP with cofilin and CAP1 enables treadmilling-like actin bundles. **a**, Schematic of the involved proteins acting on actin. **b**, Movement of large actin bundles is visible by tracking actin fluorescence over time from 30–110 min. The arrows at 30 min highlight the actin bundle’s direction of movement. **c**, Temporal-color code of the actin intensity from (**b**) shows the movement of the network. **d**, Kymographs of the actin bundle along the dotted line indicated in (**b**) further highlight the polymerizing and disassembling bundle ends for actin (red hot) decorated by zyxinΔLIM (blue). The dotted arrows indicate the velocity of polymerization and disassembly. **e**, Color coding the actin-intensity-increase (polymerization, cyan) and actin-intensity-decrease (disassembly, magenta) from 60–105 min shows the selective disassembly of small bundles and polymerization of large bundles. **f**, Quantifying velocities over time shows the rate equilibrium of disassembly and polymerization after 1 h (Mean and standard deviation of *N* = 6 − 8 per condition). **g**, Kymographs of bundle movement highlight the rates shown in (**f** ) and the polarity of growing and disassembling bundle ends. **h**, Actin bundle structure over long time scales displays an increase in bundle size around 1–2 h and disassembly of smaller bundles. **i**, Fluorescence images of zyxinΔLIM, VASP, and cofilin show the decoration of the whole bundle by the proteins. **j**, FRAP curves for zyxinΔLIM, VASP, and cofilin on actin bundles indicate the transient binding of each of the proteins. **k**, Crossover events of actin bundles (dotted arrows show direction of polymerization) indicate the preferential adhesion of zyxin-VASP to polymerizing barbed ends. At the crossover point, the non-polymerizing bundle is depleted of zyxin-VASP (white asterisks), which enables actin disassembly (red asterisk). All scale bars, 10 μm.

In contrast, in experiments lacking active disassembly, and thus no regeneration of the G-actin pool, polymerization stalls already at *t* = 15 min with bundle size limited to small widths (Fig. 1e, Extended Data Fig. 4a). We expected the transient interactions of zyxinΔLIM, VASP, and cofilin with F-actin to be essential for sustaining this treadmilling-like state of bundle movement and enabling structure-selective disassembly. To test this, we analyzed the spatial distribution and mobility of these proteins using fluorescence microscopy and FRAP. We found that actin bundles are uniformly decorated with zyxinΔLIM and VASP (Fig. 2i). In contrast to no-turnover conditions, where zyxin-VASP clusters gradually decouple from actin filaments (Extended Data Fig. 2c−e), persistent barbed-end polymerization maintains condensate localization across the entire bundle (Extended Data Fig. 3a), thereby stabilizing the structure. Cofilin, in contrast, shows a more heterogeneous distribution along the bundles, with local intensity peaks expected to correlate with pointed ends of bundles (Extended Data Fig. 4c−e). FRAP measurements reveal that zyxinΔLIM and VASP exhibit a modestly reduced diffusivity when bound to bundles, with recovery times of less than one minute (Fig. 2j). The isotropic recovery indicates diffusion on the bilayer instead of along the filaments (Extended Data Fig. 4b), while the fast recovery times highlight the transient attachment dynamics to actin in contrast with the considerably slower polymerization kinetics (*v*_*pol*_ = 0.22*μm/min*) (Fig. 2f). Cofilin shows slightly longer recovery times than zyxin or VASP, consistent with a slower unbinding rate (Fig. 2j, Extended Data Table 1). The simultaneous presence of both zyxin-VASP and cofilin on actin bundles suggests that the percolated structure of the condensates allows cofilin to access the filament. However, mechanical stabilization by the condensates likely suppresses cooperative binding and filament severing, thereby restricting cofilin and CAP1 activity to depolymerization at the pointed ends (Extended Data Fig. 4d,e).

The competition between condensate-mediated stabilization and cofilin activity becomes particularly evident at bundle crossover events (Fig.2k, Extended Data Fig. 3a−d, Supplementary Movie 2), where actively polymerizing bundles (dotted arrows) intersect with pre-existing ones. At these intersections, zyxinΔLIM-VASP condensates preferentially localize at the barbed ends of the growing bundle, resulting in a local depletion of condensate coverage on the adjacent, non-polymerizing bundle (white asterisks) (Fig.2k). This redistribution reflects the higher affinity of VASP for barbed ends relative to lateral F-actin binding. The ensuing loss of condensate protection permits unopposed cofilin activity, leading to rapid disassembly of the unprotected bundle segment (Fig.2k, Extended Data Fig.3a−d, red asterisks). These crossover events thus provide direct evidence of the dynamic competition between phase-separated stabilization and cofilin-driven disassembly.

In summary, our experimental results demonstrate that in our system, the higher-order actin bundle formation is essential for reconstituting persistent treadmilling-like dynamics. Phase-separated zyxinΔLIM-VASP condensates establish a sufficiently high local concentration of dynamic crosslinkers needed to stabilize bundles against cofilin-mediated disassembly. At the same time, the transient nature of this decoration seems to permit selective disassembly at pointed ends and of weakly crosslinked bundles. This selective disassembly and recycling of actin monomers enables the localized polymerization of highly ordered, treadmilling-like moving bundles.

### 2.3 Simulations determine optimal zyxin–VASP self-affinity driving phase separation for treadmilling-like dynamics

To investigate how zyxin-VASP phase separation regulates treadmilling-like dynamics and competes with cofilin activity, we developed an agent-based simulation framework. This approach enables systematic tuning of interaction parameters to identify the physical mechanisms governing the stability of the steady state of treadmilling-like dynamics (Fig. 3a).

**Fig. 3.**
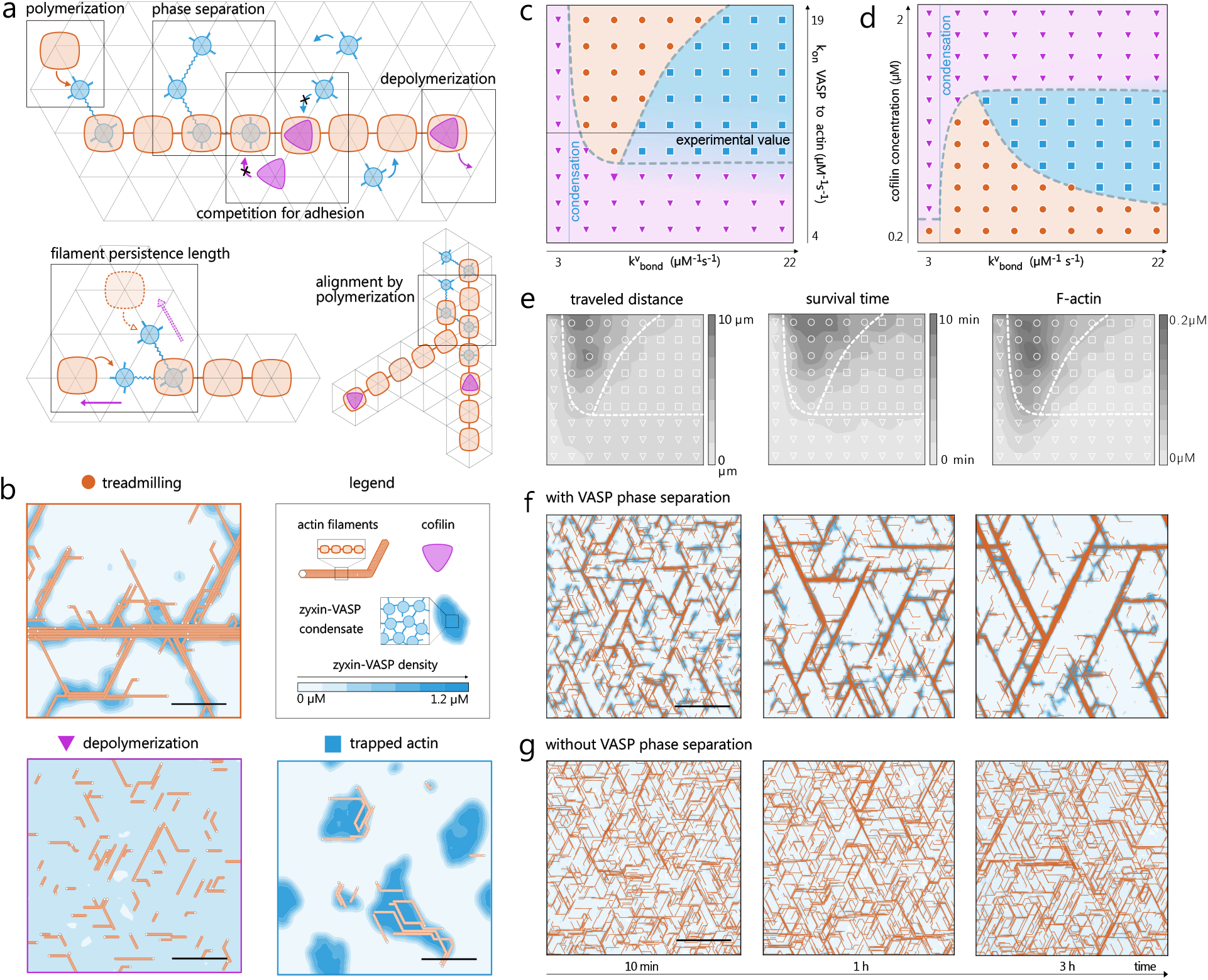
Agent-based simulations indicate treadmilling controlled by an optimum of phase separation. **a**, The simulations are conducted on a 2D hexagonal lattice with three types of agents: actin monomers (orange squares), zyxin-VASP (blue circles) and cofilin (purple triangles). Black boxes highlight the reactions included in the model. **b**, Snapshots at 1 h of three distinct simulations in the *treadmilling state* (orange circle, 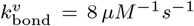 *depolymerization state* (purple triangle, 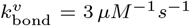, and the *trapped actin state* (blue square, 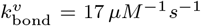 Actin filaments are shown in orange, the density of zyxin-VASP agents in blue, while the density of cofilin is not shown. The other parameter values used in these simulations are summarized in Extended Data Table 1. **c, d**, Phase diagram from our agent-based simulations, where the x-axis represents the zyxin-VASP self-affinity 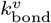. Orange, purple, and blue correspond, in that order, to the three distinct *states* shown in (**b**). In (**c**), the condensate adhesion to the filament 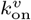 varies along the y-axis, while in (**d**), the cofilin concentration varies. The black line in (**c**) marks the experimental value of 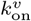 [11, 12]. The parameter values which were not varied in phase-spaces are summarized in Extended Data Table 1. **e**, Observables characterizing the three states: the average traveled distance, the average survival time of a filament before disassembly, and the amount of actin monomers in filaments (F-actin). These observables were calculated using the same parameter values as in the phase diagram (**c**), shown in the background for reference. **f**,**g**, Three snapshots showing the coarsening of filament bundles over time. In (**f** ) the zyxin-VASP complex phase-separates (parameter regime same as in **b**) so that the simulation is within the *treadmilling state*. In (**g**), the zyxin-VASP complex does *not* phase-separate 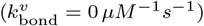 All scale bars, 15 μm.

The simulations feature three agent types: actin units, zyxin-VASP complexes, and cofilin agents. The latter represents the combined action of cofilin and CAP1. The model captures a mesoscopic representation of the underlying biochemical processes. Zyxin-VASP is modeled as a diffusive agent capable of binding and polymerizing actin bundle subunits (*polymerization*). *Phase separation* is implemented via reversible bond formation between neighboring zyxin-VASP agents, with a bond formation rate 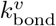 and breakage rate *k*_*break*_. The ratio 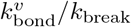 coarse-grains the effective attractive interactions between zyxin–VASP complexes in the experimental system and thereby controls condensate formation and effective surface tension(Supplementary Note 4). Since *k*_break_ is kept fixed throughout all simulations, we use the bond formation rate 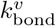 as a kinetic proxy for zyxin–VASP self-affinity, which drives phase separation in the model. *Adhesion* of zyxin-VASP along actin filaments competes with cofilin agents, which mediate *depolymerisation* and monomer recycling. This direct competition models steric exclusion and captures the stabilizing effect of VASP-mediated bundling against cofilin-induced disassembly. Additionally, because VASP crosslinking stabilizes filament bundles and prevents fragmentation, we consider only depolymerization as a mechanism for disassembly. When two zyxin-VASP agents that are mutually bonded adhere to different filaments, the complex acts as a *transient crosslinker*. Since bonds can break, the crosslinking is dynamic. To incorporate filament *persistence length* we introduce a penalty for off-axis polymerization and an additional penalty for filament crossings to account for *alignment by polymerization*, consistent with experimental observations(Fig. 1g). A concise description of the model is presented in the Methods (4), while a more detailed version is given in Supplementary Note 5.

To determine how phase separation of the zyxin-VASP complex influences depolymerization and supports treadmilling, we systematically varied key parameters in the simulations: the zyxin-VASP self affinity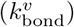 ), the adhesion of condensates to actin 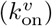 and the concentration of cofilin (Fig. 3c,d). The other rates are fixed and inferred from experimental values (Extended Data Table 1). The simulations reveal three distinct dynamical regimes: (i) a *depolymerization state*, (ii) a *treadmilling state*, and (iii) a *trapped actin state* (Fig. 3b, Supplementary Movie 3).

The *depolymerization state* arises when the zyxin-VASP self affinity 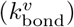 or the adhesion to actin 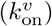 falls below a critical threshold, or at high cofilin concentrations (Fig. 3c,d). In this regime, cofilin activity dominates, leading to a net disassembly rate *v*_dis_ *> v*_pol_ and resulting in short, disordered filaments (Fig. 3b, Extended Data Fig. 8f). The effect of phase separation in mediating the competition between cofilin disassembly and zyxin-VASP crosslinking is evident in the sharp transition between the depolymerization and treadmilling states, which occurs around 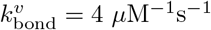.Below this threshold, the zyxin-VASP complex does not form condensates and remains uniformly distributed (Extended Data Fig. 8e, Supplementary Movie 5). Above this point, the complex undergoes phase separation into high- and low-density regions. In this regime, strong adhesion to actin causes high-density zyxin-VASP clusters to colocalize with actin bundles, while low-density regions are found in actin-depleted zones, in agreement with experimental observations (Fig. 3b,f, Extended Data Fig. 8e). The increased decoration of actin bundles by zyxin-VASP in the phase-separated regime suppresses disassembly by competing with cofilin, thereby stabilizing the bundles. This explains the sharp transition in *v*_dis_ and the onset of treadmilling (Extended Data Fig. 8f, Supplementary Note 1). Importantly, the high-density condensate value, the binodal, is set by the intrinsic phase separation properties of the zyxin-VASP complex, encoded in our model by the zyxin–VASP self-affinity, not by precise tuning of protein concentration. This enables robust treadmilling over a broad parameter range.

The *treadmilling state* observed in our simulations reproduces key features observed experimentally. The system exhibits stable treadmilling with *v*_pol_ = *v*_dis_, governed by the rate-limiting step of monomer recycling, and displays complete colocalization of zyxin-VASP agents with actin bundles (Fig. 3b). The treadmilling velocity in the simulations is found to be in good agreement with experiments (Fig. 4d, Extended Data Figure 8g, Fig. 2f). As in the experiments (Fig. 2f), the simulations show a pronounced coarsening of bundle size over time (Fig. 3f, Supplementary Movie 4). The stability and coarsening of actin bundles can be understood as a consequence of the spatial organization imposed by phase-separated condensates. Within bundles, a positive feedback loop emerges: as zyxin–VASP complexes assemble into condensates and decorate actin filaments, they recruit additional zyxin–VASP, which in turn enhances actin polymerization. This sustained polymerization promotes progressive bundle thickening and coarsening. As in experiments (Fig. 2e), outside the bundles, filaments lose VASP coverage and are rapidly disassembled. Thus, phase separation of zyxin-VASP both stabilizes treadmilling and spatially organizes these bundles, leading to their coarsening.

**Fig. 4.**
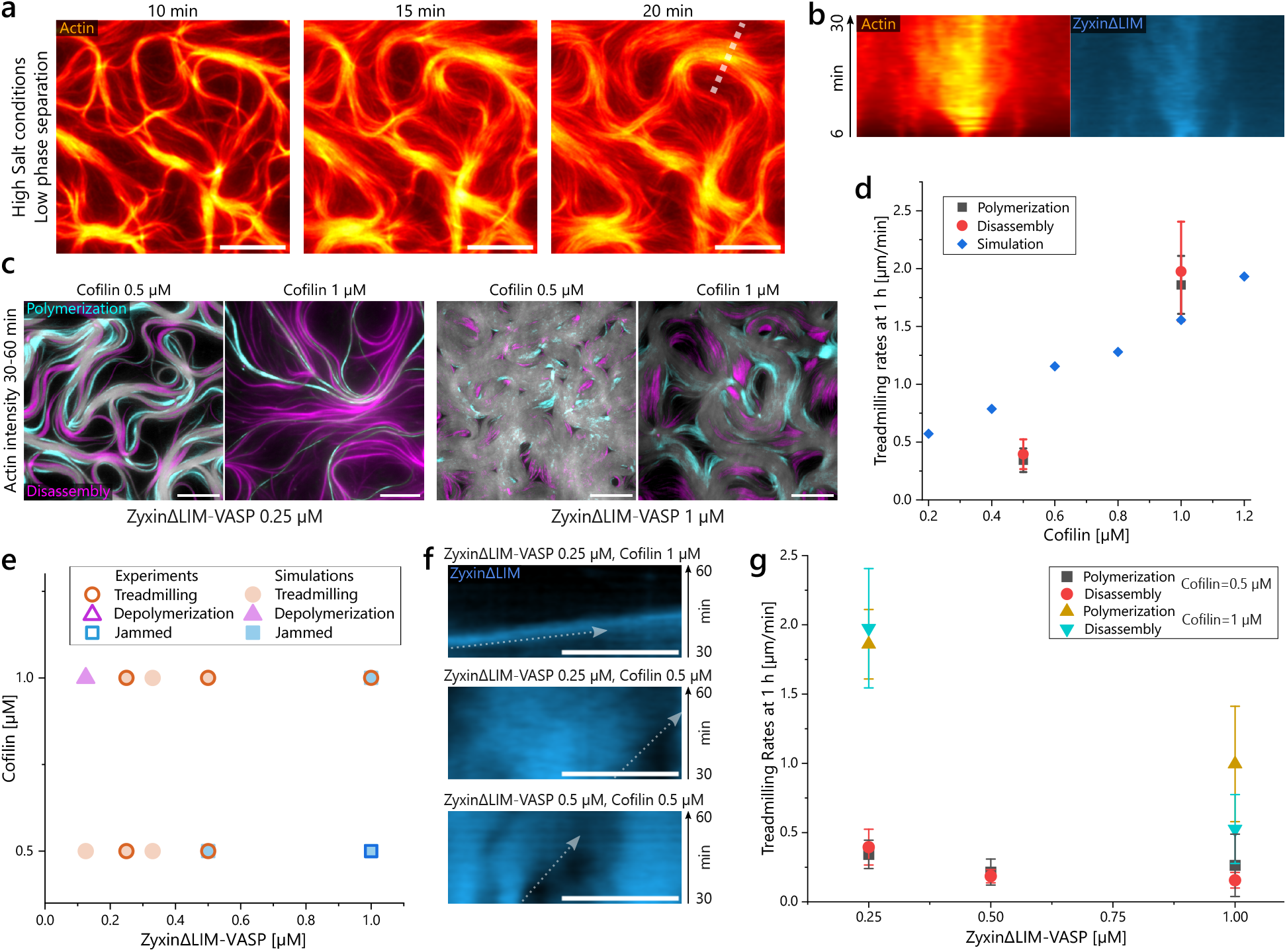
Competition of the phase-separated zyxin-VASP and cofilin tunes treadmilling-like actin bundle velocity. **a**, Actin bundles formed by zyxinΔLIM-VASP (*C* = 0.5 μM) are disassembled under high salt concentration (100 mM KCl). **b**, The kymograph indicated by the dotted line in (**a**) displays the bundle fragmentation and splaying over time. **c**, Actin fluorescence intensity difference showing the actin polymerization (cyan) and disassembly (magenta) between *t* = 30− 60 min overlaid over the actin network at *t* =60 min highlights the degree of network rearrangement by turnover. **d**, Simulated treadmilling-like dynamics and corresponding experimentally determined velocities increase in dependence of cofilin concentration. **e**, Assessed state of actin networks for different concentrations of zyxin-VASP and cofilin between experiments and simulations. **f**, Experimental velocities shown by the slope in kymographs of bundles decorated with zyxinΔLIM highlight the higher velocities at *C*_*cofilin*_ = 1*μM* . **g**, Polymerization and disassembly rates of dynamic bundles at *t* = 1 h for different zyxinΔLIM-VASP and cofilin concentrations (mean and standard deviation of *N* = 4 − 13 per rate). All scale bars, 10 μm.

These results show that liquid-liquid phase separation of zyxin-VASP not only stabilizes treadmilling but also imposes spatial structure, driving bundle coarsening over time. In contrast, in the absence of phase separation, treadmilling occurs only at significantly reduced cofilin concentrations (Fig. 3d), since no mechanism stabilizing the local value of VASP density is present in this case. As a consequence, a drastically reduced range of concentrations exhibits treadmilling. Under these conditions, bundles exhibit minimal coarsening, and disordered filaments persist outside the bundle cores (Fig. 3g).

The *trapped actin state* emerges at high values of the zyxin-VASP self-affinity 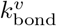 (Fig. 3c,d). This regime is characterized by arrested bundle movement, accompanied by a marked reduction in both *traveled distance* and F-actin content (Fig. 3e). In fact, a high self-affinity 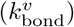 increases the effective surface tension of the condensates (Supplementary Note 4, Supplementary Movie 5), leading to spatial confinement of actin filaments and restricting polymerization to the interior of the droplets. Filaments that polymerize beyond the condensate edge lose VASP decoration (Extended Data Fig. 8b) and are rapidly depolymerized, while bundles at the interface remain trapped and fail to elongate (Fig. 3b). A detailed analysis of how filament stiffness, condensate surface tension, and polymerization velocity determine the boundary of this regime is provided in Supplementary Note 2, 4, and Extended Data Fig. 8a–c,h. Taken together, the agent-based simulations identify an optimal range of zyxin-VASP self-affinity, driving phase separation, that supports stable treadmilling by balancing polymerization and depolymerisation dynamics. In contrast, deviations from this optimal range lead either to disassembly due to insufficient condensation or to actin trapping due to overly strong cohesion.

### 2.4 Modifying zyxin-VASP self-affinity tunes competition with cofilin

The simulations predict that the emergence of a stable *treadmilling state* instead of a *depolymerization state* requires a threshold concentration of zyxin-VASP on the actin bundles, which is set by phase separation, driven by the self-affinity, of the protein complex. To experimentally validate the simulated *depolymerisation state* at low zyxin-VASP self-affinity 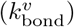 we increased the salt concentration from 50 mM to 100 mM KCl, weakening the electrostatic interactions within zyxinΔLIM-VASP complexes and with F-actin (Extended Data Fig. 1g,h). Actin polymerization and initial network formation (*t* = 10 min) are unaffected at this salt level (Fig. 4a). However, upon onset of cofilin and CAP1 activity on ADP-F-actin, the zyxinΔLIM-VASP condensates are displaced from the bundles, resulting in a splitting of the bundles into single filaments and complete disassembly (Fig. 4a,b). Under these conditions, bundle movement is abolished. These results confirm that cohesive interactions within the zyxin-VASP condensates are essential for maintaining filament bundling, which, in turn, is required to sustain competition with cofilin and preserve a treadmilling-like state.

An effective intermediate self-affinity of phase-separated zyxinΔLIM-VASP supports robust treadmilling-like dynamics across a broad range of concentrations of cofilin and zyxinΔLIM-VASP (Fig. 4c, Extended Data Fig. 5a,b). Only at high concentrations (*C*_zyxinΔLIM_ = *C*_VASP_ = 1*μ*M), treadmilling-like movement is limited (Fig. 4c,f,g). Under these conditions, the bilayer becomes densely covered with nematically aligned actin filaments, resulting in steric hindrance that limits movement and restricts polymerization to short bursts emerging from +1*/*2 nematic defects (Extended Data Fig. 6c− e, Supplementary Movie 7). A similar transition is observed in simulations at comparable concentrations (Fig. 4e, Extended Data Fig. 8d). Consistent with simulation results, increasing the cofilin concentration experimentally leads to a marked acceleration of treadmilling-like dynamics. Doubling the cofilin concentration to *C*_cofilin_ = 1*μ*M results in a four-fold increase in velocity of bundle dynamics and monomer concentration (Fig. 4d,g, Extended Data Fig. 5b,c,e, Supplementary Movie 6). This increase in turnover is accompanied by pronounced bundle coarsening (Extended Data Fig. 5c, Supplementary Movie 6). A similar trend of enhanced actin dynamics and coarsening at increased cofilin concentration is also observed at high concentrations of zyxinΔLIM and VASP (Fig. 4c,g, Extended Data Fig. 6a,b,e, Supplementary Movie 7). These results confirm that increased monomer recycling through cofilin and CAP1 boosts both filament turnover and the growth of large, mobile actin bundles.

The agent-based model also predicts a transition to a *trapped actin state* at high amounts of self-affinity of zyxin-VASP, where treadmilling-like movement becomes arrested due to an excessive condensate surface tension (induced by a high. 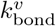) To experimentally validate this regime, we used zyxinFL-VASP condensates, which exhibit additional multivalent interactions and thus increased self-affinity as a complex compared to zyxinΔLIM-VASP (Fig. 5a). The LIM domains in zyxinFL enhance both self-association and binding to actin filaments, increasing condensate stability.

**Fig. 5.**
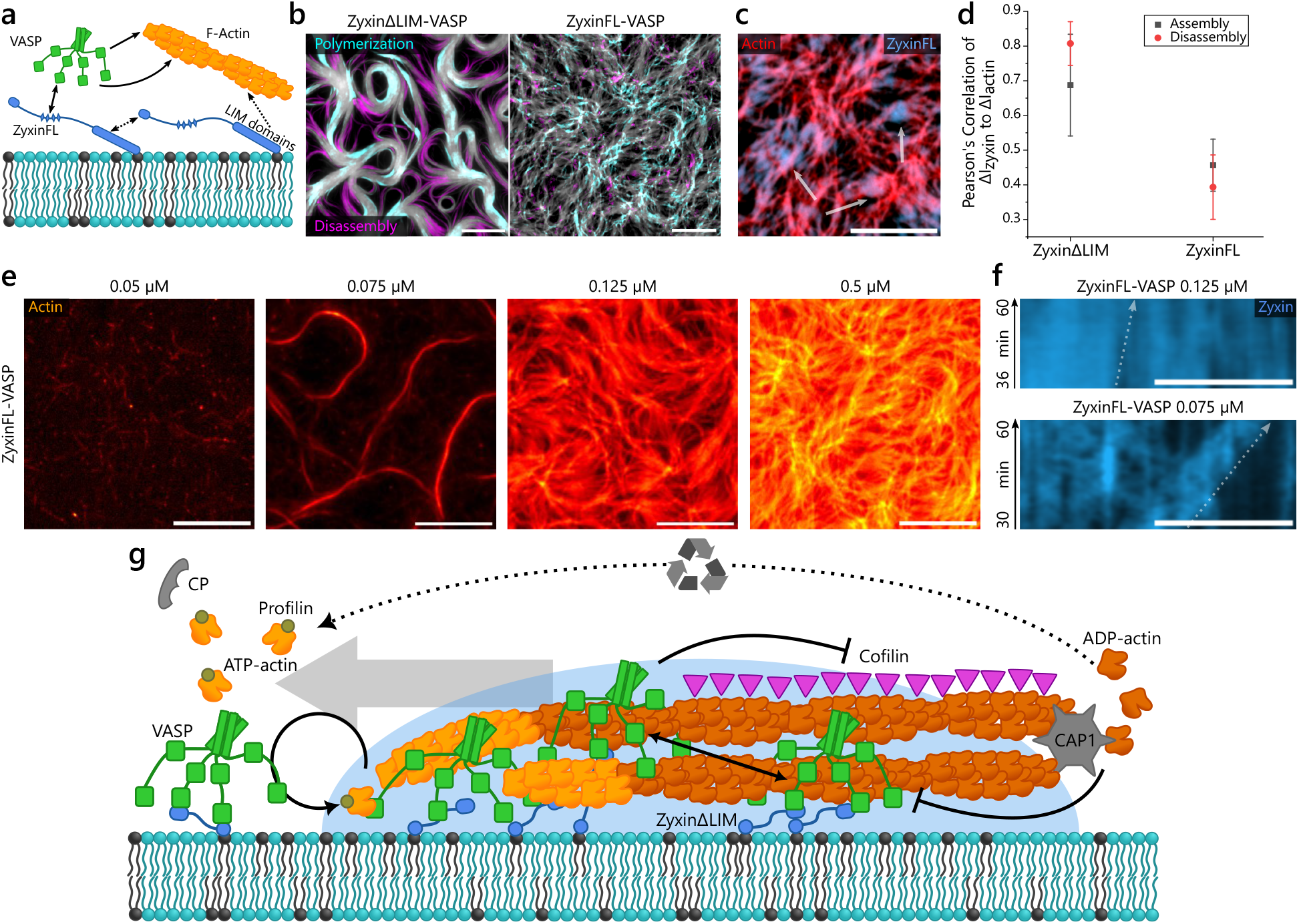
Increased interactions of zyxin-VASP condensates severely limit treadmilling-like movement. **a**, Schematic displaying the interactions of zyxin and VASP with actin. Additional interactions of zyxinFL are indicated by dotted arrows. These limit the condensate’s fluidity and the competition with cofilin. **b**, Actin-intensity-increase (polymerization, cyan) and actin-intensity-decrease (disassembly, magenta) from t=30-60 min shows the disordered polymerization of zyxinFL-VASP compared to zyxinΔLIM-VASP. **c**, Fluorescence images of zyxinFL bound to actin (*t* = 66 min) show actin-turnover-independent formation of large clustered structures (white arrows) over time. **d**, Correlation of positive (associated with actin assembly) and negative (associated with disassembly) intensity changes from *t* = 33 − 66 min between actin and zyxin (mean and standard deviation of *N* = 3 − 4 per condition). **e**, Actin fluorescence at 1 h shows no actin at *C* = 0.05*µM* zyxinFL/VASP concentration, treadmilling-like actin bundles at *C* = 0.075*µM* and disordered actin filaments at concentrations higher than *C* = 0.125*µM* zyxinFL and VASP. **f**, Kymograph of actin bundle movement as shown by the decorating zyxinFL fluorescence. Movement ceases at *C* = 0.125*µM* zyxinFL and VASP. **g**, Summary of the observed treadmilling-like mechanism. Zyxin and VASP condensates polymerize and accumulate on actin bundles. In a positive feedback loop, the polymerizing barbed end recruits more zyxin-VASP and actin by monomer funneling. The bundle stability, based on filament size and a stabilizing-effect through the zyxin-VASP self-affinity, inhibits disassembly by cofilin. Cofilin and CAP1, in turn, displace the condensates from the depolymerizing pointed end and recycle the actin monomers. All scale bars, 10 µm.

Following the formation of an initially disordered actin network during polymerization (Fig. 5b, Extended Data Fig. 7a− c), the strong self-affinity of zyxinFL-VASP condensates promote slow formation of large, spherical domains on the network *t* = 66 min (Fig. 5c, Extended Data Fig. 7a). While the experimental network initially resembles an isotropic, polymerizing system rather than the *trapped actin state* seen in simulations, the emergence of binodal-like spherical condensates driven by a high self-affinity proceeds in agreement with the simulated predictions (Fig. 3b, Extended Data Fig. 8a− c, Supplementary Movie 8). The high surface tension of zyxinFL-VASP condensates opposes their stable adhesion to actin filaments, resulting in reduced colocalization between condensates and actin. This is quantified by the correlation of fluorescence intensity changes (Δ*I*) between zyxin and actin, which is significantly lower for zyxinFL-VASP than for zyxinΔLIM-VASP (Fig. 5d, Extended Data Fig. 7b). This shows a mechanical decoupling of condensate dynamics from the actin dynamics. In addition, the increased binding affinity of zyxinFL to F-actin limits cofilin activity, impairing disassembly and thus suppressing the monomer concentration for polymerization. As in the simulations, this results in a pronounced reduction or complete arrest of actin bundle motility (Fig. 5f, Extended Data Fig. 7d, 8a−c). At the low concentration of *C*_zyxinFL_ = *C*_VASP_ = 75 nM, below the threshold for full condensate gelation (Extended Data Fig. 1c), the mobility of zyxinFL-VASP clusters is sufficient to allow bundle formation and controlled cofilin activity, which are prerequisites for treadmilling-like movement (Fig. 5e,f, Extended Data Fig. 7e,f,h, Supplementary Movie 8). Even under these conditions, however, the still elevated surface tension leads to gaps in bundle decoration (asterisks), which trigger local filament fragmentation (Extended Data Fig. 7e−g). Thus, the treadmilling-like bundle movement, limited by the surface tension for zyxinFL-VASP clusters, contrasts the robust and continuous dynamics observed across a broad range of concentrations for zyxinΔLIM-VASP.

By integrating agent-based simulations with in vitro measurements, we show that the phase separation properties of zyxin-VASP are critical for enabling robust treadmilling-like movement of actin bundles (Fig. 5g). Transient crosslinking by condensates, which stiffens bundles in a size-dependent manner, allows them to resist cofilin-mediated disassembly. This competition is governed by a threshold concentration of zyxin-VASP on actin bundles, maintained by their high-density phase. At the same time, condensate fluidity permits depolymerization at the pointed ends. A key emergent mechanism is the spatial funneling of actin monomers towards condensate-stabilized bundles, where active polymerization sustains the localization of the condensates. This self-organized feedback promotes spatially confined polymerization, leading to bundle coarsening and long-lived actin bundle motility. These dynamics crucially require an intermediate strength of self-affinity of zyxin-VASP. A weak self-affinity strength prevents phase-separation and actin is insufficiently bundled. Whereas if the self-affinity of the condensates is too high, the condensate surface tension decouples the condensate-localization from actin dynamics, arresting turnover. Thus, stable treadmilling-like movement emerges from a finely tuned balance between self-affinity of condensates, filament binding, and protein mobility.

## 3 Discussion

Actin filaments require tightly regulated assembly and disassembly dynamics, orchestrated by a complex network of actin-binding proteins, to support diverse cellular functions [1, 9]. This regulatory complexity has so far hindered the reconstitution of persistent, motile, treadmilling-based actin assemblies in vitro [10]. Experimental systems have so far been limited to reconstituting actin network turnover instead of filament treadmilling. This was done by nucleating actin filaments at solid supports using either branched actin networks [23–26] or linear filament networks [27, 28]. In these setups, the balance of growth and turnover is achieved through spatial separation of nucleation at the surface and distally located stochastic fragmentation by cofilin [25, 26]. Thus, modulating nucleation or severing rates tunes the network length without intrinsic coupling of these processes. In contrast, here, we observe a robust system of treadmilling-like motion based on the reciprocal coupling of condensates polymerizing and decorating the filaments and the disassembly by cofilin. This results in the emergence of highly-ordered actin structures, including aligned bundles and vortices. While the ordered structure of filaments initially arises from alignment interactions consistent with results from propelled filaments [4, 5, 46–48], in our system, the higher-order organization is formed as a prerequisite for the biochemical coupling of turnover rates. The structure-selective disassembly and monomer funneling provide a self-reinforcing mechanism promoting actin domain growth, unlike treadmilling systems with fixed or independently tuned rates [2, 49–51]. Strikingly, the formation of large vortices is stabilized without requiring intrinsically chiral assembly of filaments [3, 52]. While some redistribution of monomers has previously been observed in bundles stabilized by conventional crosslinkers in combination with active disassembly, such systems typically arrest in non-treadmilling states [28–30, 32]. In contrast, here, the zyxin-VASP condensates allow continuous monomer recycling for actin bundle motility. The condensates further promote the formation of spatio-temporally persistent, highly-ordered actin structures that retain a dynamic nature by enabling monomer funneling and locally reinforced, processive polymerization.

Liquid-liquid phase separation of proteins has been widely implicated with the localized nucleation of cytoskeletal filaments [37, 38, 45, 53–56]. We expand on this known role in cytoskeletal regulation by presenting a mechanism where the molecular condensates additionally influence filament dynamics by decorating actin bundles and altering the activity of other binding proteins. This could present a broad model for the regulation of filament turnover for a wide range of known filament-wetting condensates [57].

The selective assembly of highly ordered actin bundles from a disordered network in our reconstituted system offers a mechanistic model for the formation of bundled actin in cells such as filopodia and microspikes, which emerge from the disordered lamellipodium [58, 59]. These bundled actin arrays treadmill at rates comparable to those observed in our experiments and simulations [60–63], and exhibit similar nematic alignment and coarsening during bundle growth [58–60]. Additionally, their formation depends on membrane-bound actin dynamics [60, 63, 64] and often features or depends on an elevated VASP concentration [58, 61, 63, 65]. In particular, some turnover-based actin waves have been shown to be based on VASP [64] or the VASP-zyxin complex [63]. Since VASP can form phase-separated condensates with a range of binding partners [35, 36, 45, 66], the proposed regulatory mechanism, based on the material properties of the multivalent protein condensates, may represent a broader role in the organization of actin-rich structures.

## 4 Methods

### 4.1 Protein purification

Actin was purified from rabbit skeletal muscle acetone powder based on a previously published protocol [67]. No rabbits were directly involved in this study. G-actin was stored in 2mM Tris/HCl pH 8.0, 0.2mM ATP, 0.2mM CaCl_2_, 0.2mM DTT, and 0.005% NaN_3_. Fluorescent actin was prepared by labeling exposed lysines with an Atto-488 NHS-ester dye (Jena Biosciences, FP-201-488). Freshly dialyzed actin was used within a week. For differently labeled actin, Atto-532 NHS-ester (Jena Biosciences, FP-201-532) was used.

The other proteins were recombinantly expressed in *Escherichia coli* BL21 CodonPlus DE3-RIPL (Agilent Technologies, 230280). The heterodimeric capping protein (CP) was expressed in a pRFSDuet-1 vector (Novagen, 71341). Here, the mouse *α*1- and human *β*2 subunits of CP were used. Mouse profilin-2a was expressed in a pGEX-6P-2 vector (Cytiva, GE28-9546-50). CP and profilin were purified the same way as previously described [37].

Human cofilin-1 and as well as the cofilin-mCherry fusion protein was also expressed using a pGEX-6P-2 vector. Mouse cyclase-associated protein-1 (CAP1) was expressed using a pET-28b(+) vector (Novagen, 69864). Cofilin and CAP1 were purified according to a previous study [27].

A human VASP construct containing the G-actin binding domain of *Dictyostelium discoideum* in a pET-28a(+)-TEV vector (GenScript) was used [11]. VASP was purified as described before [37] with an additional step to cleave off the His-tag. To achieve this, the size exclusion column was run with 30mM Hepes, 150mM KCl, 0.5mM EDTA, and 5% (vol/vol) glycerol. The His-tag was cleaved off overnight at 10 °C using TEV-protease (Serva, 36401). After addition of 20mM imidazole to the sample, the tag was removed by incubating the proteins with cOmplete His-Tag purification resin (Roche, 5893801001) for 30 min and collecting the flow through. For labeling, VASP was incubated with an Alexa fluor-532 maleimide dye (Invitrogen, A10255) as previously described [37]. The His-tag-free VASP was subsequently concentrated and flash-frozen in liquid nitrogen. The point mutations L229A, L230A, I233A, and L241A in the G-actin binding domain and R273A, R274A, E275A, and K276A in the F-actin binding domain were inserted into VASP using the QuikChange Multi Site-Directed Mutagenesis Kit (Agilent, 200514) to create a mutant incapable of binding to actin.

Human zyxin constructs were expressed and purified in a pET-28a(+)-TEV vector. Zyxin full-length (zyxinFL) and zyxin without the LIM domains (amino acids 1–372) (zyxinΔLIM) were C-terminally fused to mCherry to produce labeled proteins. An uncleavable C-terminal His-Tag was kept for the zyxin constructs, to capture the cellular localization of zyxin via the C-terminal LIM domains. After the main *e. coli* culture reached an optical density at 600 nm of 0.6-0.8, 2mM betaine-HCl was added to the culture and left to shake for 1 h at 23 °C. Expression of zyxin was induced using 0.5mM isopropyl-*β*-D-thiogalactopyranoside (IPTG) and shaking at 2 °C for 12 h. The bacteria were harvested and lysed by freeze–thaw and French pressing in lysis buffer containing 50mM Tris/HCl pH 7.5, 300mM NaCl, 1mM EGTA, 20mM imidazole, 1mM DTT,10 % (vol/vol) glycerol, 40 µg/ml lysozyme, one tablet of cOmplete protease inhibitor (Roche, 4693132001), and 2 U/ml Benzonase (Sigma, E1014). After centrifugation, the supernatant was filtered and incubated to equilibrated cOmplete His-Tag purification resin (Roche, 5893801001). The His-Tag purification was done following standard procedures. The proteins were eluted using 50mM Tris/HCl pH 7.5, 300mM NaCl, 1mM EGTA, 500mM imidazole, 1mM DTT, 10 % (vol/vol) glycerol, concentrated and the buffer exchanged to size exclusion buffer (30mM HEPES/HCl pH 7.5, 200mM KCl, 5 % (vol/vol) glycerol, and 1mM DTT) supplemented with 0.5mM EDTA using a NAP-25 column (Cytiva, GE17-0852-02). The N-terminal His-Tag was then cleaved using TEV-Protease (Serva, 36401) and incubation at 10 °C overnight. Afterwards, a HiLoad 16/600 Superdex 200 size-exclusion column (Cytiva, 11397490) was run with size-exclusion buffer. Zyxin was then concentrated, aliquoted, and flash-frozen.

### 4.2 Supported lipid bilayers

The supported lipid bilayers were created by incubating small unilamellar vesicles on hydrophilic glass. The glass slides for the supported lipid bilayers (SLB) were first cleaned with 3 M NaOH and sonication for 30 min. The slides were further washed with double-distilled *H*_2_*O* (*ddH*_2_*O*) and etched with piranha acid for 5 min. Piranha acid was prepared by carefully mixing 100 ml *H*_2_*SO*_4_ with 50 ml *H*_2_*O*_2_ under stirring and incubating in a water bath at 60 °C. After the treatment with piranha acid, the coverslips were then extensively washed and then stored in *ddH*_2_*O*. The glass slides as basis for the chamber formation were cleaned by sonication in 2 % Hellmanex III (Hellma, Z805939-1EA) for 30 min. They were further washed with *ddH*_2_*O* and stored in 100 % ethanol.

For the small unilamellar vesicles, a 20mM lipid-in-chloroform mix of 87.45 % egg-PC (L-*α*-phosphatidylcholine from chicken egg) (Sigma, P3556), 10 % DOGS-Ni-NTA (1,2-dioleoyl-sn-glycero-3-[(N-(5-amino-1-carboxypentyl)iminodiacetic acid)succinyl]) (nickel salt) (Avanti Polar Lipids, 790404), 2.5 % DSPE-PEG2000 (1,2-distearoyl-sn-glycero-3-phosphoethanolamine-N-[methoxy(polyethylene glycol)-2000]) (ammonium salt) (Avanti Polar Lipids, 880120), and 0.05 % DSPE-Cy5 (1,2-distearoyl-sn-glycero-3-phosphoethanolamine-N-[Cyanine 5]) (Avanti Polar Lipids, 810345C) was prepared. 50 µl of the lipid mix were dried under a nitrogen stream and dessicated under vacuum for at least 3 h. The lipids were then resuspended to 1mM in phosphate-buffered saline pH 7.4 (PBS) by vortexing for 30 s and sonication in a water bath for 30 min. The vesicles were further passed 10 times through a 100 nm pore size Whatman Nuclepore membrane (Sigma, WHA111105) using a lipid extruder (Avanti Polar Lipids, 610000-1EA) and stored at 4 °C. The small unilamellar vesicles, coverslips, and glass slides were all used within one week.

### 4.3 Treadmilling-like experiments

The experiments were done in KMEI buffer (10mM imidazole/HCl pH 7.5, 50mM KCl, 1mM MgCl_2_, 1mM EGTA, 1mM DTT). The concentrations for all the proteins were calculated as the monomeric proteins, except for CP, for which the heterodimeric complex was used. All buffers used for the SLB experiments were degassed for at least 30 min directly before use. For every experiment a fresh supported lipid bilayer was prepared. A flow chamber of around 10 µl volume was made from three layers of parafilm, the prepared coverslips, and the glass slides. The lipid mix was diluted to 0.2mM in PBS and incubated to the flow chamber for 20 min. Afterwards, the flow chamber was washed with 1.5 ml of PBS. For the actin polymerization experiments, we first incubated the His-tagged zyxin construct to the SLB for 10 min in PBS with 1mM DTT. The chamber was washed with 1.5 ml of PBS (phosphate-buffered saline) and 100 µl KMEI buffer. The buffer for incubation and wash of zyxinFL additionally contained 2.5 µM ZnCl_2_ to ensure functional LIM domains. An equimolar concentration of VASP was incubated to zyxin in KMEI buffer for 10 min. The channel was washed with 150 µl of KMEI buffer supplemented with 0.1 mg/ml BSA. The polymerization buffer consisted of 2 µM actin with 12.5 % labeled with atto-488, 10 µM profilin, 50 nM CP, and 2mM ATP in KMEI buffer. For experiments with active turnover, 0.5 µM cofilin, 0.125 µM CAP1, and an ATP-regeneration system consisting of 10mM creatine phosphate (Roche, 10621714001) and 40 U/ml creatine phosphokinase (Sigma-Adrich, C3755) was added to the polymerization mix. The ATP-regeneration constituents were prepared as 30x stock solutions in 89mM Tris, 89mM *H*_3_*BO*_3_ pH 8.3. As bleach protection agents, we added catalase at 1700 U/ml (Sigma-Aldirch, C40), glucose-oxidase at 26 U/ml (Sigma-Aldrich, G2133) or pyranose-oxidase at 10 U/ml (Sigma-Aldrich, P4234), and 36mM glucose. Actin and glucose were added to the mix directly before the start of the measurement. After addition of actin, the flow chamber was quickly sealed with vacuum grease and the imaging started signifying time point t=0 min.

### 4.4 Image acquisition

Fluorescence images were taken on a Leica DMi8 with a 100x objective (numerical aperture 1.47) with oil immersion and an ORCA-Flash 4.0 CMOS camera (Hamamatsu, C13440-20CU). Cy5-lipids, VASP, and zyxin were imaged using epifluorescence. Actin and cofilin were imaged using total internal reflection fluorescence with a Leica Infinity TIRF HP module. FRAP measurements were done using a Leica Infinity scanner and bleaching a circle of 5 µm in diameter.

### 4.5 Data analysis

Image analysis was performed using FIJI-ImageJ [68]. Fluorescence intensities were quantified on raw image data with subtracted background intensity. Fluorescence recovery after photobleaching (FRAP) was analyzed by first applying a drift correction [69]. The intensity was adjusted by correcting for bleaching using a correction factor calculated from an unbleached region. The intensity was further normalized using the prebleach intensity and the first frame after bleaching. The recovery curves were fitted using the double exponential function *I*(*t*) = *A*_1_ *×* (1− *e*^−*t/τ*1^ ) +*A*_2_ *×* (1− *e*^−*t/τ*2^ ). The mobile fraction results from *A*_1_ +*A*_2_. The half-time *t*_1*/*2_ of reformation of droplets after arrested polymerization was fitted to the zyxin-to-actin-bundles fluorescence intensity ratio using the logistic function 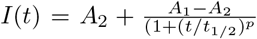. The actual *t*_1*/*2_ is then obtained by subtracting the time of the initial polymerization (*t* = 7.4 *min*).

The fluorescence intensity of bundles was determined as the mean intensity over 5 bundles. For zyxin, the intensity ratio was determined by the intensity on the bundle compared to the intensity adjacent and void of bundles. The actin intensity ratio *I*_*on*_*/I*_*off*_ was calculated from the 5 largest bundles of the networks. The bundles were straightened, and the intensity over 1 µm diameter on the bundles was divided by the intensity of a 1 µm wide area directly adjacent to the bundles. The angles of binary interactions during polymerization were determined by the direction of polymerization over time and manually measured using the angle tool in FIJI-ImageJ for the incoming angle before collision and the outgoing angle.

The velocities of the treadmilling-like dynamics were determined as the slopes of intensity increase or decrease from kymographs on the bundles. The kymographs were created as the average over 5 pixels of the diameter using the KymoResliceWide plugin [70]. The polymerization and disassembly velocity was then determined as the mean with standard deviations of the angle for intensity increase or decrease in the kymographs. Local cofilin intensity peaks were calculated by applying a rolling ball correction with a radius of 1 pixel. The VASP-mediated polymerization on-rate *k*_*on*_ was fitted to the equation *v*_*pol*_ = *k*_*on*_*C*_*actin*_ −*k*_*off*_ using the initial single filament polymerization velocity *v*_*pol*_ in subunits per second, the known initial actin concentration of *C*_*actin*_ = 2*µM*, and an off-rate of *k*_*off*_ = 1 *±*3 *s*^−1^ [71]. The steady-state actin monomer concentrations could be approximated using the same equation using the steady-state *v*_*pol*_ and the determined on-rate of *k*_*on*_ = 33.8*±* 0.2*µM* ^−1^*s*^−1^.

For the correlation analysis of intensity difference between zyxin and actin, the positive and negative intensity differences between two time points with a 30 min time span were calculated. Pearson’s correlation coefficient was then calculated separately for the positive (polymerization) and negative (disassembly) intensity difference images between zyxin and actin using the Coloc2 plugin in FIJI-ImageJ. Particle image velocimetry was performed between two time points t=30-60 min using the iterative PIV plugin [72].

Bundle width was determined from images that were segmented using ilastik [73]. The width was calculated as local diameter along the distance ridge of the bundle using the local thickness plugin in FIJI ImageJ [74].

### 4.6 Numerical Model

The model captures a coarse-grained version of the minimal reaction set inferred from the known biochemistry to reproduce the observed phenomenology. Agent-based simulations were performed on a 2D lattice with three types of mesoscale agents: actin (100 monomers per agent), zyxin–VASP complexes (100 VASP and 100 zyxin molecules per agent), and cofilin (100 cofilin molecules per agent). A triangular lattice was chosen to provide more realistic turning angles, consistent with filament stiffness, and to better approximate isotropic space compared to a square lattice. Boundary conditions are periodic to avoid finite-size effects. At each time step, agents are updated sequentially. Each agent can execute one reaction per time step (*dt* = 0.001), and reactions are chosen probabilistically, according to their reaction rate (Supplementary Note 5). Note that diffusion, implemented as hopping to nearest-neighbors, is modeled as a separate process from on-site reactions.

VASP agents, confined to the membrane, can form up to four reversible bonds (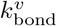 and 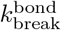), mimicking tetrameric interactions and enabling clustering via phase separation. Agents diffuse by 2D hopping between neighboring sites or along filaments via a diffuse-and-capture mechanism, where a VASP agent reaching the barbed end cannot hop back. This leads to a higher local density of VASP at filament barbed ends [75], mimicking the preferential adhesion of VASP to barbed ends observed experimentally. VASP agents can also attach to 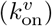 or detach from 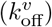 actin filaments. Diffusion is restricted to moves that do not stretch existing bonds, effectively reducing the mobility of clustered agents relative to individual ones, as observed experimentally. To mimic steric repulsion between VASP proteins, a maximum of five agents per site is enforced. Filament-associated cofilin agents can detach 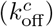, diffuse (modeled as instantaneous relocation due to 3D bulk diffusion), and reattach 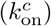. Cooperative binding of cofilin and barbed-end tracking by VASP are omitted due to the mesoscopic resolution of the model. VASP and cofilin compete for binding to the same filament site to reflect their mutually exclusive activities.

Actin agents represent filamentous (F-)actin explicitly on the lattice, while monomeric (G-)actin are treated as a homogeneous bulk concentration. Polymerization (*k*_pol_) occurs at filament tips in the presence of colocalized VASP agents and consumed G-actin. Directional growth and energy penalties for filament crossing (*µ*_cross_) and off-axis growth (*µ*_bend_) are included to reflect experimental observations of alignment. Disassembly (*k*_dis_) at filament ends is driven by cofilin, replenishing the G-actin pool. Filaments shorter than two monomers are considered “dead” and cannot depolymerize further, though they may re-elongate. The total number of actin monomers (in either G- or F-actin states) is conserved throughout the simulation. Simulations are initialized with randomly placed filaments of length 7. All rate conversions to physical units are provided in Supplementary Note 3, and final parameter values are listed in Extended Data Table 1.

## 5 Data Availability

Source data and raw data necessary for the reproduction of the image analysis will be publicly available on Zenodo (https://doi.org/10.5281/zenodo.17226587).

## 6 Code Availability

A minimal working version of the code used for the agent-based simulations in this study is publicly available on Zenodo (https://doi.org/10.5281/zenodo.17277200).

## 7 Acknowledgments

This work was supported by the ERC via the project PoInt (810104) and CellGeom (101097810). We further acknowledge financial support from the Deutsche Forschungsgemeinschaft (DFG, German Research Foundation) through the Excellence Cluster ORIGINS under Germany’s Excellence Strategy (EXC-2094-390783311). EF acknowledges additional support from the Chan–Zuckerberg Initiative (CZI). We thank Monika Rusp-Post, Karin Vogt and Gabriele Chmel for the preparation of actin and the support during the purification of CP and profilin. We thank Philip Bleicher for establishing the purification protocols for cofilin and CAP1. We thank Prof. Jan Faix (Cytoskeleton Dynamics, Medizinische Hochschule Hannover) for gifting the plasmid for VASP.

## 8 Author Contributions

T.N. designed, performed, and analyzed the experiments. T.N. purified the proteins. B.N., D.T., and M.S. designed the simulations. B.N. and D.T. performed and analyzed the simulations. T.N., B.N., D.T., E.F., and A.R.B. wrote and revised the manuscript.

