## Extended Data Figures and Supplemental Notes for "Phase separation determines treadmilling-like movement of actin bundles"

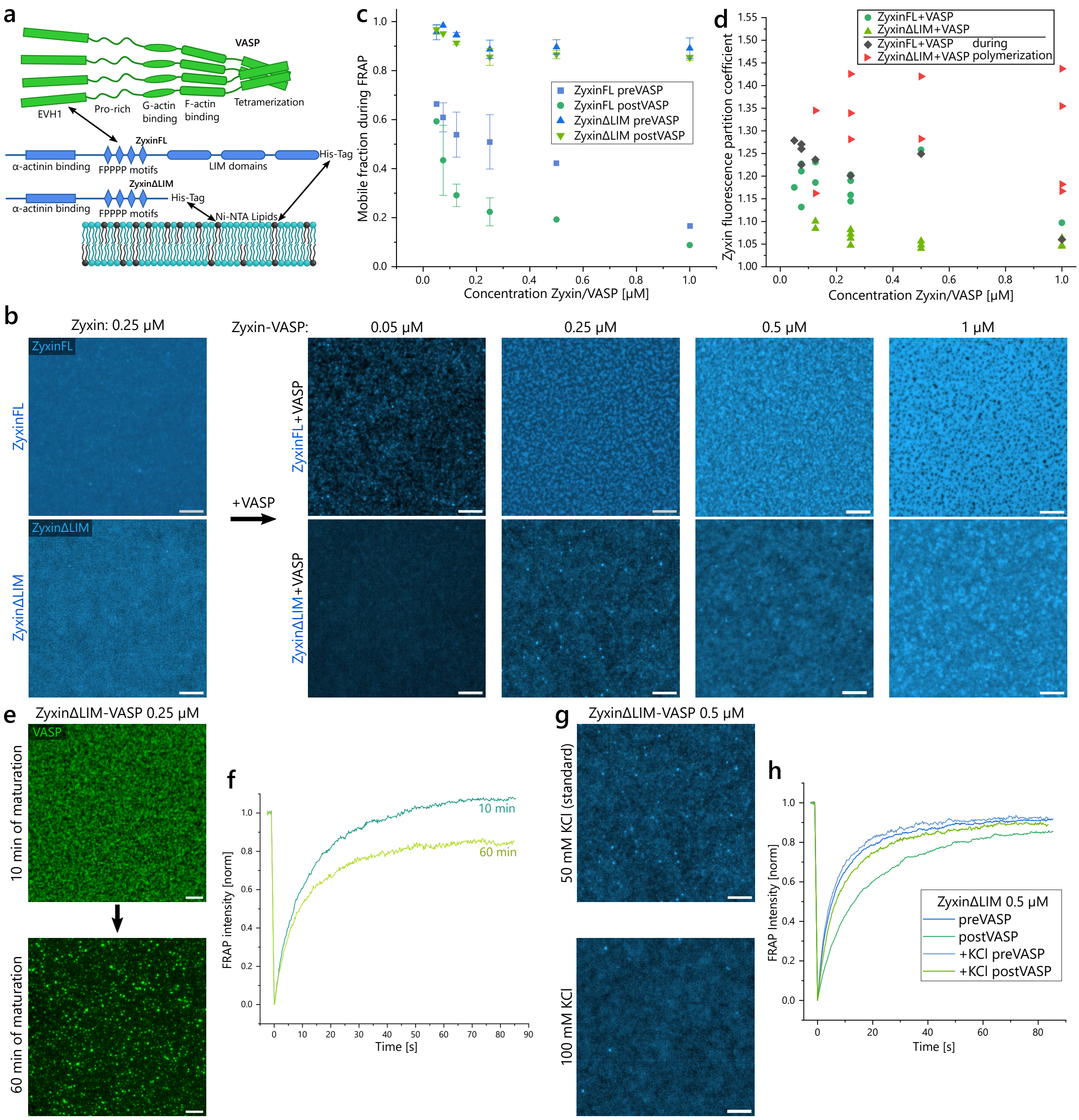


**Extended Data Fig. 1 Mobility of the phase separation of zyxin and VASP. a,** Interactions between VASP's EVH1 domain and the FPPPP motifs on zyxin enable phase separation on the bilayer. **b,** The mobile fraction of FRAP decreases with VASP concentration. ZyxinFL in comparison to zyxinΔLIM shows a VASP-independent decrease at increasing concentrations. Mean and standard deviation from N(total)=87 experiments. **c,** Addition of equal concentration of VASP to zyxin (mCherry-labeled) shows condensation of zyxin after 10 min of incubation. The size of condensates increases with concentration of the proteins. **d,** Partition coefficient of zyxin fluorescence shows higher partitioning for zyxinFL-VASP before addition of actin and enhanced partitioning for zyxinΔLIM-VASP during actin polymerization ($t = 9$ min). **e,** Fluorescent VASP (Alexa 532) on zyxinΔLIM displays a clustered structure after 10 min that further condenses with longer maturation. **f,** FRAP experiments corresponding to (**e**) highlight the lower mobility of the increased condensation after 60 min. **g,** Additional 50 mM KCl in the buffer (to 100 mM KCl total) reduces the number of zyxinΔLIM-VASP clusters compared to clusters formed at 50 mM KCl. **h,** FRAP intensity for zyxinΔLIM before and after addition of VASP with or without high KCl concentration shows that additional KCl prevents the VASP-associated decrease in fluidity. All scale bars, 5 μm.

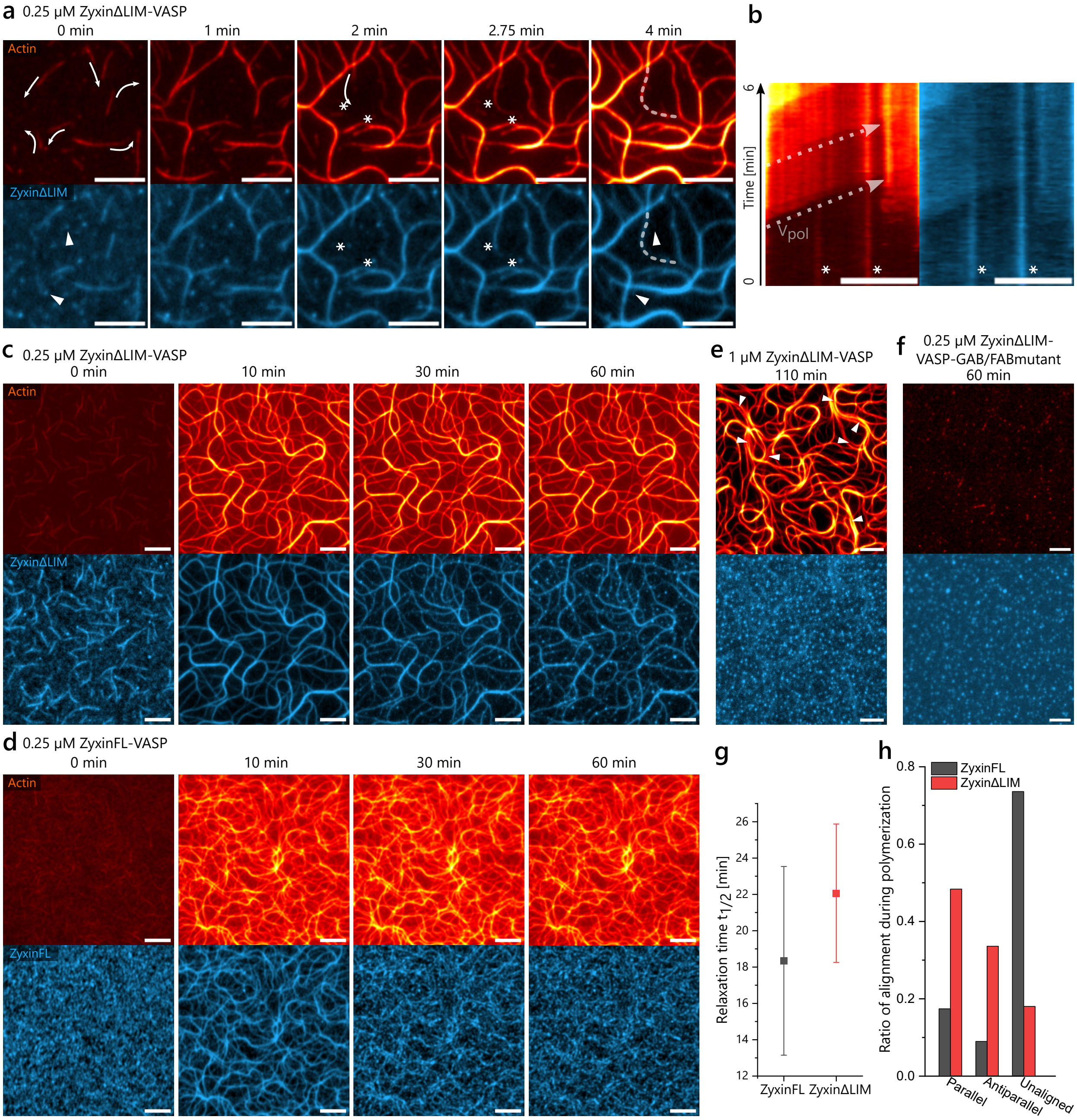


**Extended Data Fig. 2 Condensate colocalization during actin polymerization. a,** Early polymerization acquisition shows the redistribution of zyxinΔLIM-VASP droplets along polymerizing actin filaments. Direction of polymerization is indicated by arrows, also showing the alignment of filaments during polymerization. Droplets being spread along polymerizing filaments are indicated by asterisks, and droplets that redistribute by an Ostwald-ripening-like process are shown with triangles. **b,** Kymograph along the dotted line in (**a**) highlights the distribution of droplets (asterisks) along the direction of actin polymerization (dotted arrow). **c, d,** Time-dependent decoration of actin with zyxinΔLIM-VASP (**c**) and zyxinFL-VASP (**d**) condensates ($C_{zyxin} = C_{VASP} = 0.25\,\mu M$) during long-term polymerization shows complete decoration during active actin polymerization ($t = 10$min). The condensates form separate filament-associated droplets after polymerization stalls out (t=60 min). **e,** Larger bundles formed by $C_{zyxin\Delta LIM} = C_{VASP} = 1\,\mu M$ visualize the splitting of bundles (triangles) after the condensates lose decoration of the bundle and form independent droplets ($t = 110$min). **f,** VASP, mutated to prevent G-actin- and F-actin-binding, bound to zyxinΔLIM shows that without VASP's activity no actin polymerization takes place and no actin-polymerization-dependent deformation of the condensates is visible. **g,** Quantification of relaxation half-time $t_{1/2}$ of zyxin fluorescence on actin bundles after the colocalization peak from Fig. 1e shows a faster relaxation towards actin-independent droplets for zyxinFL-VASP condensates (mean and standard deviation of $N_{zyxinFL} = 5$ and $N_{zyxin\Delta LIM} = 10$). **h,** Quantification of filament alignment during initial polymerization from Fig. 1g. All scale bars, 5 µm.

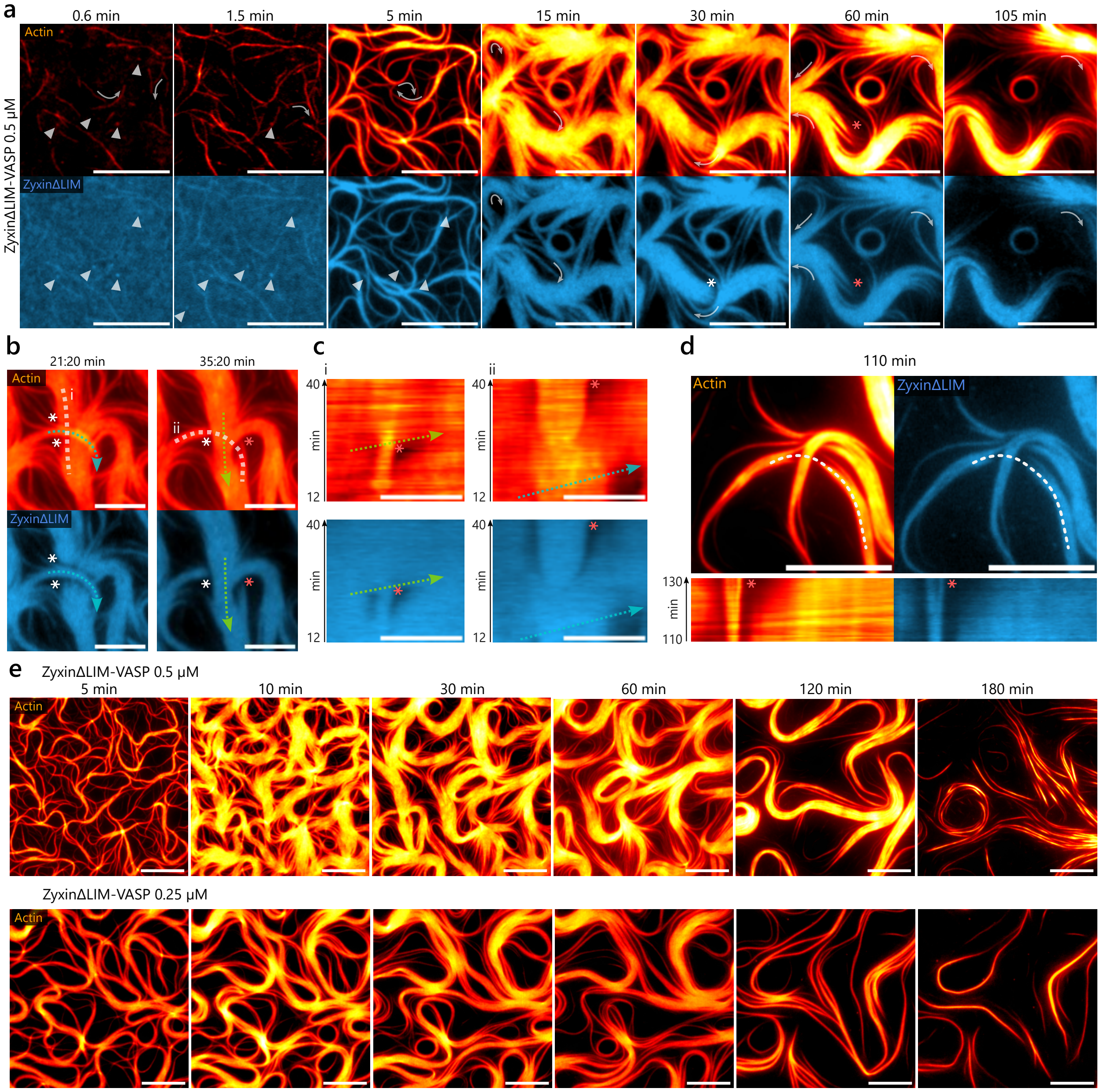


**Extended Data Fig. 3 Actin structure formation by treadmilling-like dynamics.** **a,** Actin is polymerized by zyxinΔLIM-VASP condensates ($C = 0.5\,\mu M$) into large bundles with treadmilling-like movement. Triangles indicate condensate clusters merging with the actin-bundle-associated condensates. The direction of polymerization during treadmilling-like movement is indicated with arrows. Bundle crossover events lead to displacement of zyxinΔLIM-VASP condensates as shown by asterisks, which can results in actin depolymerization (red asterisks). **b,** Corresponding actin fluorescence images of the bundle crossover shown in Fig. 2k shows a persistent actin bundle upon short zyxinΔLIM-VASP depletion (white asterisk), whereas long zyxinΔLIM-VASP depletion causes actin bundle disassembly (red asterisk). **c**, Kymographs of actin and zyxinΔLIM fluorescence along the dotted lines (**i, ii**) in (**b**) show the crossover-induced zyxin depletion and actin disassembly. Colored arrows (cyan, green) highlight the sequential bundle polymerization events. **d,** Kymographs over time along the dotted line of a bundle crossover event at a later time point emphasizes directional actin disassembly (red asterisk) originating from filament fragmentation induced by the zyxinΔLIM-VASP condensate depletion. **e**, Actin bundles formed by ZyxinΔLIM-VASP condensates at $C = 0.25 - 0.5\,\mu M$ show a time-dependent coarsening of bundle width. All scale bars, 10 µm.

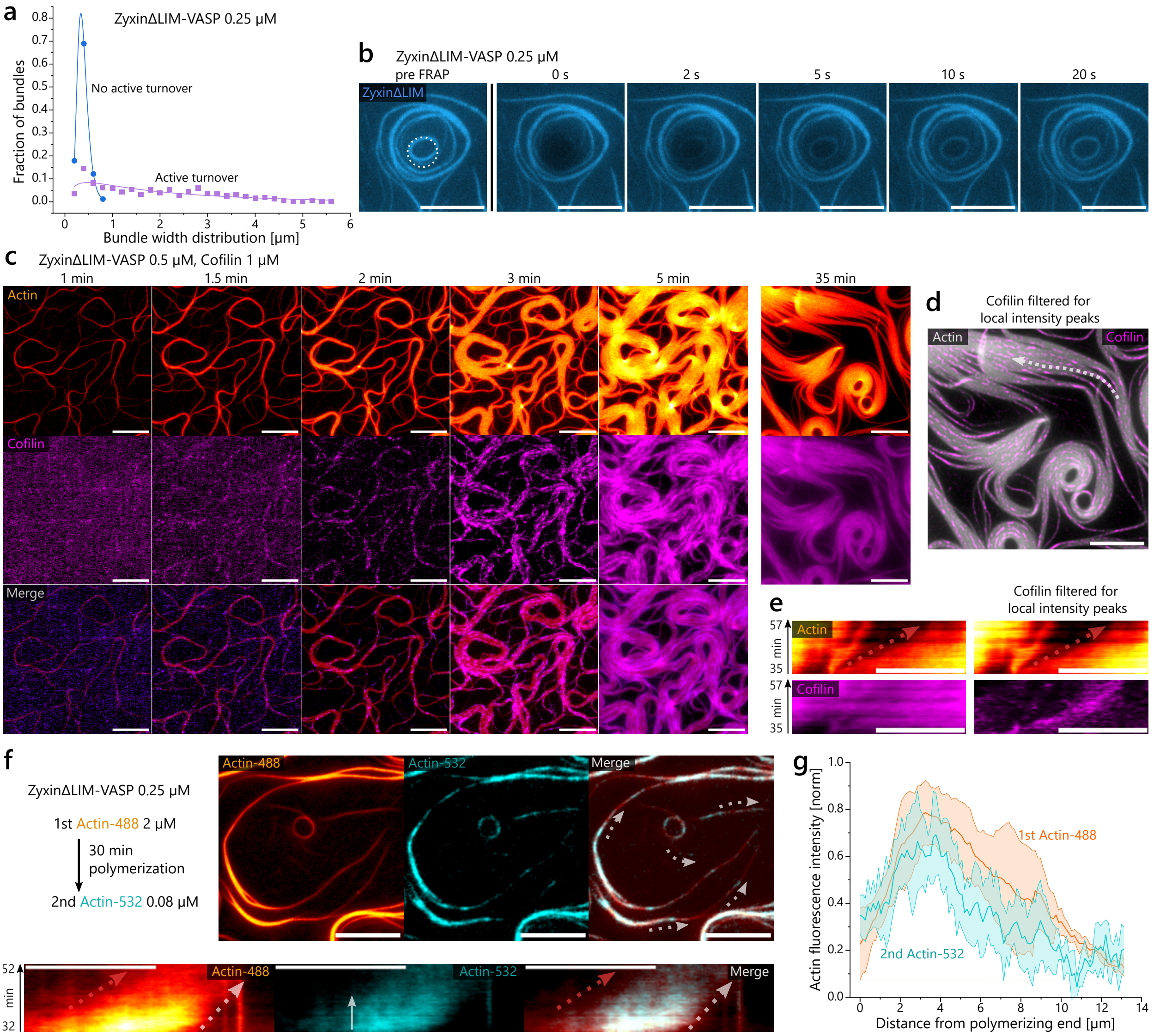


**Extended Data Fig. 4 Distribution of barbed ends and pointed ends throughout bundles. a,** Quantified distribution of bundle width at $t = 30\,min$ shows the drastic increase in width by addition of active turnover (cofilin, CAP1) in the system. The histogram is fitted with a log-normal distribution. **b,** FRAP experiment of zyxinΔLIM-VASP ($C = 0.25\,\mu M$) bound to an actin bundle vortex displays diffusion in 2D on the bilayer. **c,** Cofilin fluorescence shows shows small clusters localizing to actin fluorescence after 1.5 min to almost complete actin decoration. Decoration is not completely homogeneous with higher intensity clusters on the actin bundles. Cofilin contrast is adjusted for $t = 3$ min onward to account for the drastic increase in fluorescence intensity. **d,** Filtering for local high intensity cofilin clusters (rolling ball subtraction with 1 pixel radius) visualizes the aligned structure within the bundles and the high amount of bundle pointed ends. **e,** Kymograph along the dotted line in (**d**) for unfiltered cofilin fluorescence and filtered fluorescence (right) shows colocalization of local intensity cofilin clusters to depolymerizing bundle ends (red dotted arrow). **f,** Subsequent addition of actin, labeled with Atto-532, to bundles polymerized with actin labeled with Atto-488 for 30 min shows initial incorporation of new actin in isolated clusters at small bundles and along the whole bundle for larger bundles. A kymograph highlights the progression in time of incorporated actin-532 along a small bundle. The gray arrow indicates an edge of incorporated actin-532 in the bundle that temporally persists until disassembly at the pointed end of the bundle. Grey dotted arrows indicate direction of polymerization, while the red dotted arrow shows disassembly. **g,** Fluorescence localization of new actin-532 and existing actin-488 directly after addition of actin-532 in small bundles shows preferential localization at the polymerizing barbed ends of the bundles (mean and standard deviation of $N = 5$). All scale bars, 10 μm.

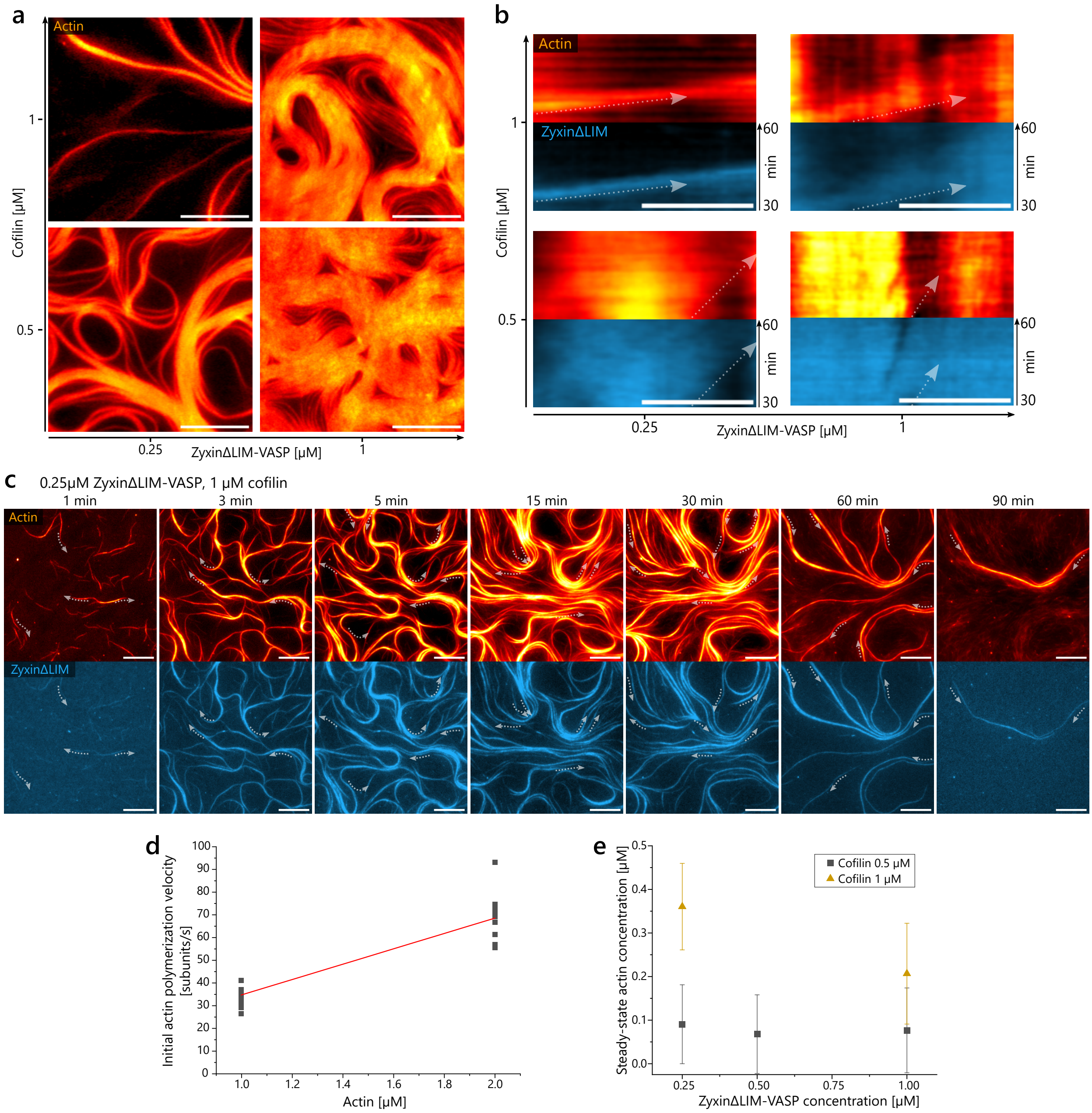


**Extended Data Fig. 5 Accelerated treadmilling-like dynamics at high cofilin concentration. a,** Actin fluorescence images of the bundle structure formed after 1 h by different concentrations of cofilin and zyxinΔLIM-VASP. CAP1 is present at a ratio of 1:4 of cofilin concentration. **b,** Kymograph of labeled actin and zyxinΔLIM shows bundle movement from t=30-60 min. **c,** Fluorescence images show fast actin treadmilling-like movement and accelerated bundle coarsening at $C_{zyxin\Delta LIM-VASP} = 0.25\,\mu M$ and $C_{cofilin} = 1\,\mu M$. Dotted arrows indicated direction of polymerization of bundles. **d,** Quantified initial actin polymerization velocities at different starting actin concentrations show the VASP-specific apparent polymerization on-rate. A fit (red line) using the published off-rate at VASP-bound barbed ends of $k_{off} = 1 \pm 3\,s^{-1}$ [1] results in an apparent on-rate of $k_{on} = 33.8 \pm 0.2\,\mu M^{-1}s^{-1}$. $N = 12$ per concentration. **e,** Actin monomer concentrations calculated from steady state polymerization velocities at $t = 1$ h using $k_{off}$ and the fitted $k_{on}$ from (**d**) at different concentrations of cofilin and zyxinΔLIM-VASP. All scale bars, 10 μm.

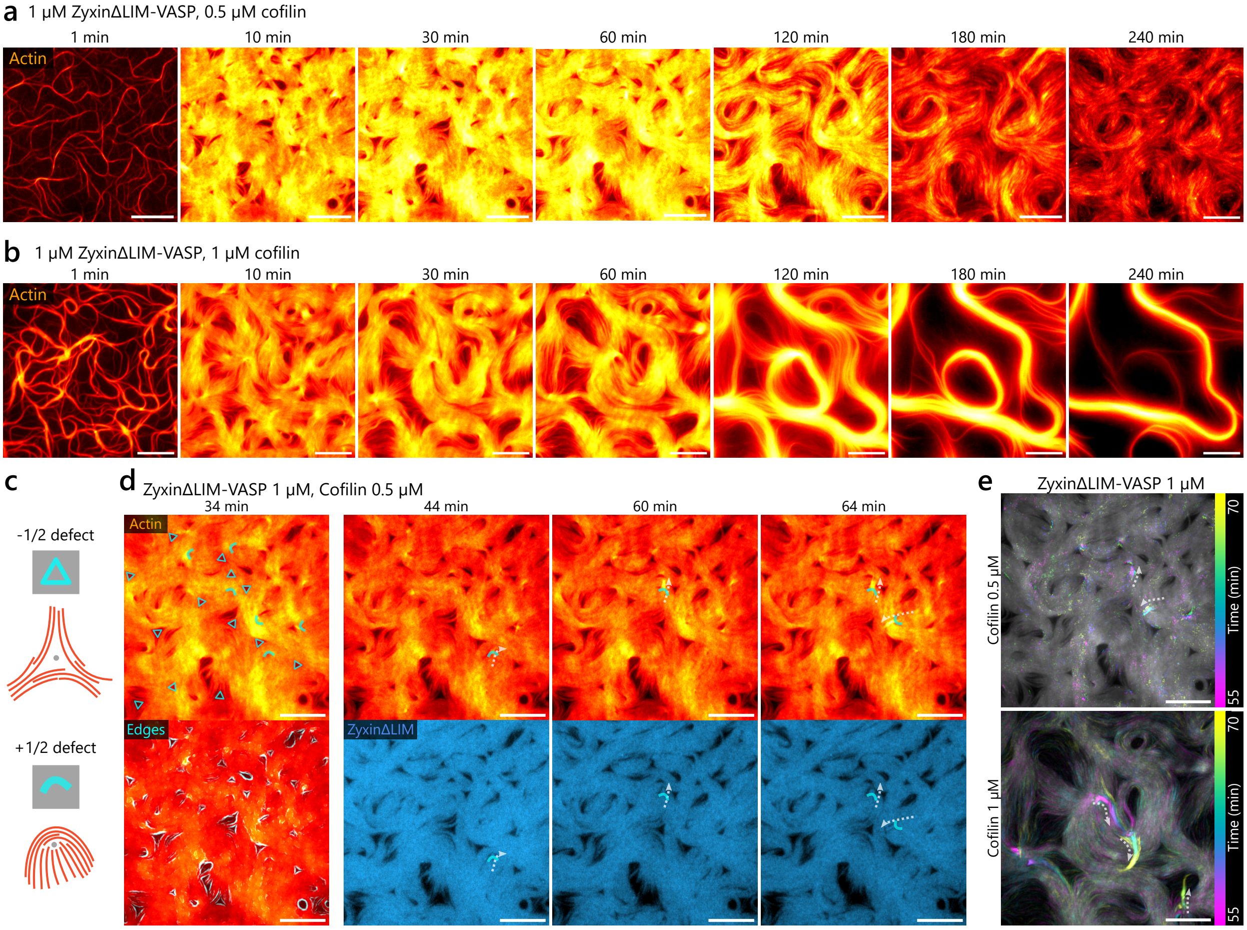


**Extended Data Fig. 6 Limited treadmilling-like dynamics at high concentrations of zyxinΔLIM-VASP. a,b,** At $C_{zyxin\Delta LIM-VASP} = 1\,\mu M$, a bilayer decorating bundle network is formed with bundles jammed at nematic defects limiting treadmilling-like dynamics. With $C_{cofilin} = 1\,\mu M$ (**b**), over time pronounced bundle coarsening is observable. **c,** Schematic representation of the filament structure leading to -1/2 and +1/2 defects. **d,** Edge detection (cyan) of actin intensity allows visualization of +1/2 (curves) and -1/2 nematic defects (triangles) in the actin decoration formed by zyxinΔLIM-VASP ($C = 1\,\mu M$) and cofilin $C = 0.5\,\mu M$. Short polymerization bursts of accumulated bundles can be seen from +1/2 defects (dotted arrows). **e,** Temporal color code of actin intensity of the polymerization shown in (**d**) and more pronounced polymerization bursts from nematic defects at increased cofilin ($C = 1\,\mu M$). All scale bars, 10 µm.

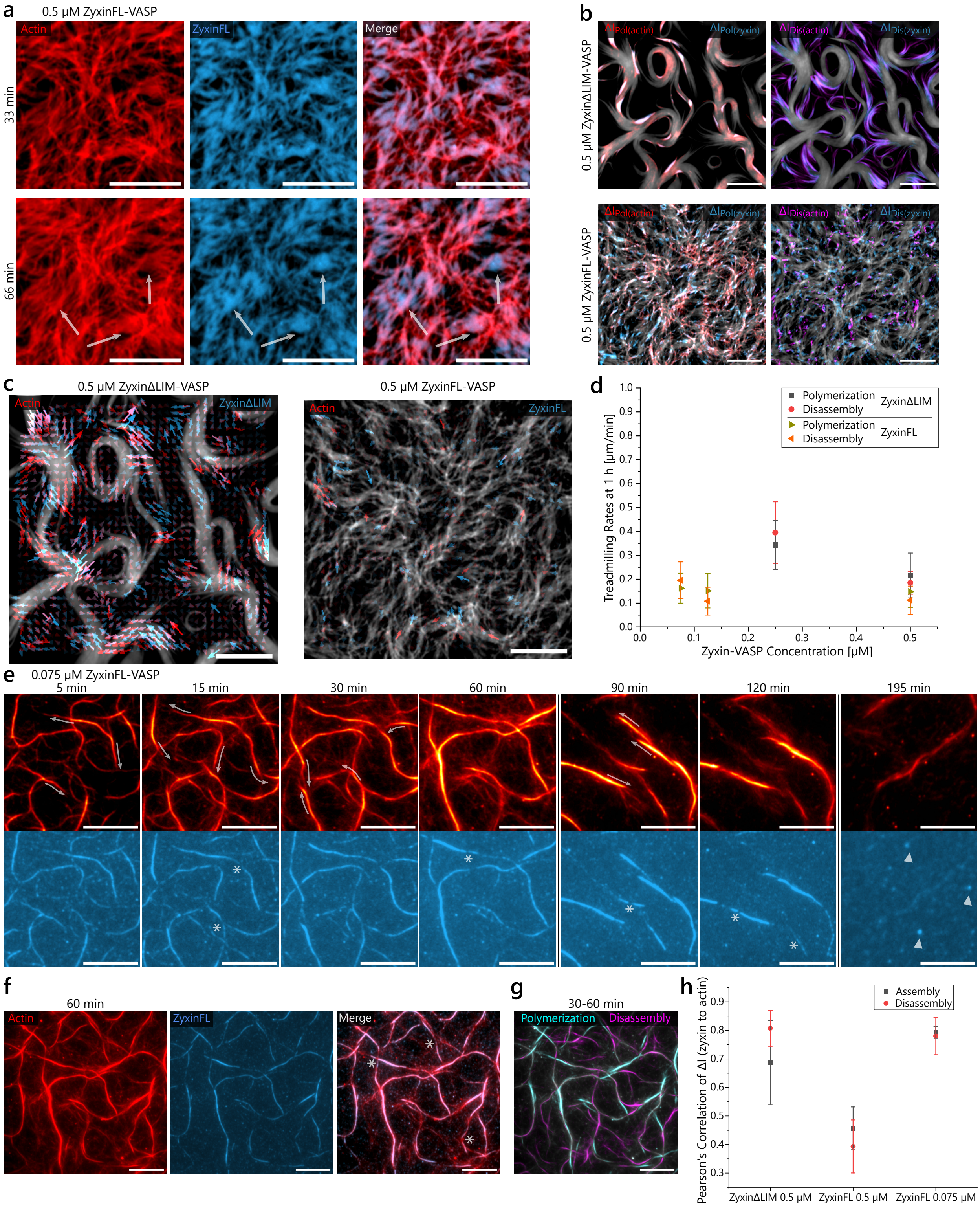


**Extended Data Fig. 7 ZyxinFL-VASP condensates inhibit actin bundling and treadmilling-like movement. a,** Fluorescence images over time show little directional movement of actin bundles. ZyxinFL fluorescence is not homogeneously colocalized to actin and forms large domains decoupled from the actin intensity indicated by the arrows. **b,** Colocalization of the intensity differences from t=30-60 min associated with polymerization ($\Delta I_{Pol}$) and disassembly ($\Delta I_{Dis}$) for actin and zyxin are overlaid over the actin bundle network. Intensity differences are colocalized for zyxinΔLIM-VASP and actin and show highly coordinated movement, whereas zyxinFL-VASP is less colocalized to actin and the actin intensity differences are disordered. **c,** Particle image velocimetry analysis overlaid over the actin network highlights the coupled movement of actin with zyxinΔLIM-VASP and the disordered and arrested dynamics with zyxinFL-VASP. **d,** Quantification of detectable bundle movement from kymographs shows inhibited disassembly and movement for zyxinFL-VASP above a concentration of $C_{zyxinFL-VASP} > 0.125\,\mu M$. (Mean and standard deviation from N=8-20 bundles per velocity from 1-3 experiments per condition) **e,** At a concentration of $C_{zyxinFL-VASP} = 0.075\,\mu M$ movement of isolated bundles is visible. Polymerization direction is shown with arrows and lines between time points indicate a different field-of-view. High surface tension of zyxinFL-VASP condensates can lead to gaps in the homogeneous decoration of the actin bundles (asterisks). This is highlighted by the overlay in **f**. Displaced zyxinFL-VASP condensates form round clusters on the bilayer (triangles). **g,** Intensity difference images of actin overlaid with the actin bundles shows the directionality of treadmilling-like movement. **h,** Quantification of the correlation of the intensity change of zyxinFL and actin shows a high colocalization for $C_{zyxinFL-VASP} = 0.075\,\mu M$ (Mean and standard deviation of N=3-4 from 2 experiments per condition). All scale bars, 10 µm.

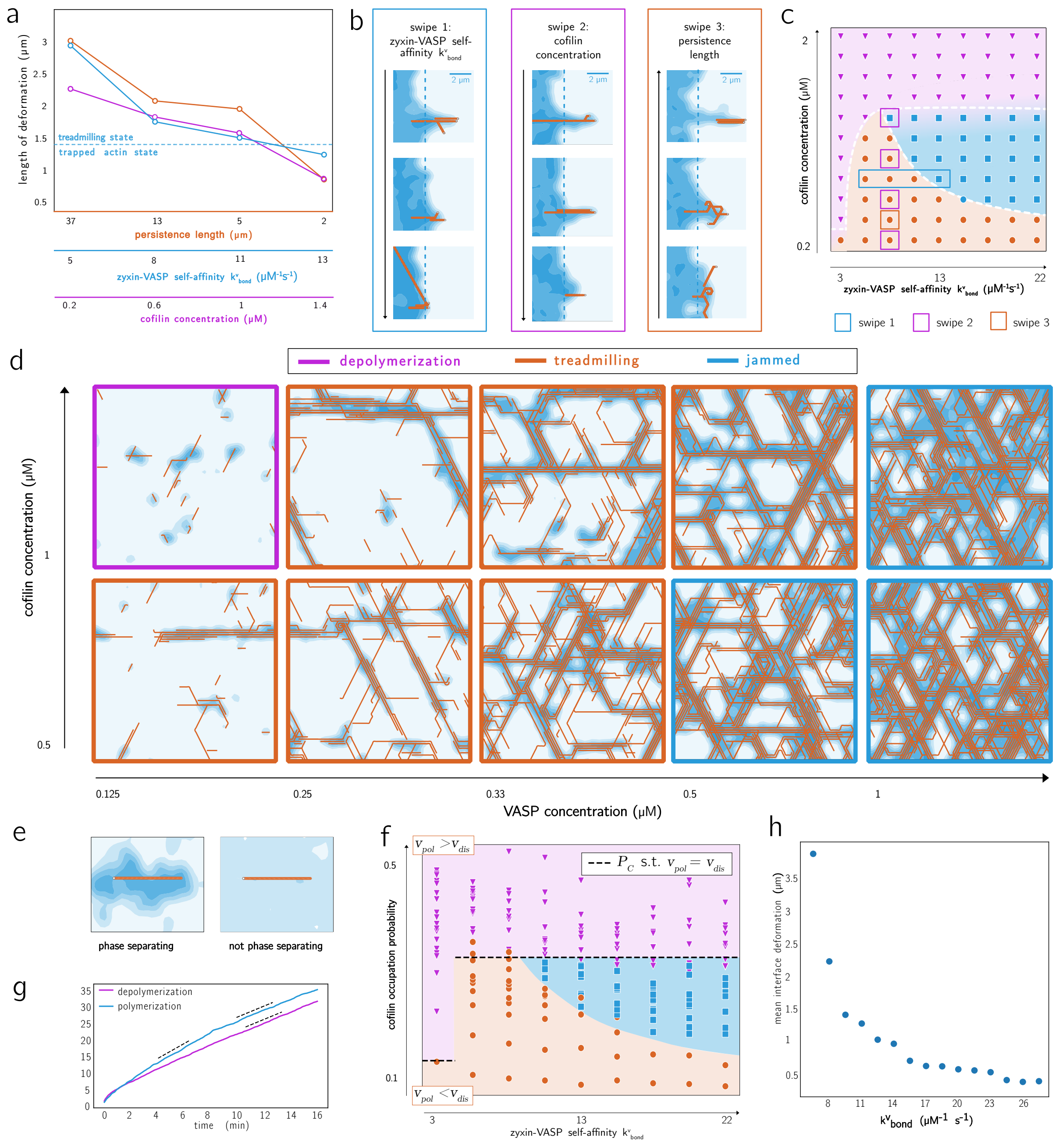

**Extended Data Fig. 8 Additional data on numerical results. a,** Mean elongation after 9 minutes of a flat VASP interface induced by bundles of three filaments placed perpendicular to the VASP interface. We vary the zyxin-VASP self-affinity $k^v_{\text{bond}}$ (blue), the cofilin concentration (purple), and the filament persistence length, tuned by varying varying the off-axis growth penalty (22). The mean elongation was obtained by averaging over 10 simulations per parameter set. **b,** Three representative series of how the interface deformation varies when changing the three parameters of zyxin-VASP self-affinity ($k^v_{\text{bond}}$), cofilin concentration and filament stiffness. The dashed line in blue indicates the initial position of the interface. **c,** The phase diagram indicating where the simulations parameters in **a** and **b** lie. **d,** Phase diagram obtained by varying cofilin and VASP concentrations, the same quantities varied in Fig 4e. At high VASP concentrations, a new *jammed state* emerges, in line with the experimental observations (Extended Data Fig. 6). Parameters use are the same as in Fig. 2, in the *treadmilling state*. System size $20\,\mu m$. **e,** Snapshots of two simulations consisting of a single static filament, one in the phase-separating regime $k^v_{\text{bond}} = 5\,\mu M^{-1}s^{-1}$ and one in the non-phase-separating regime $k^v_{\text{bond}} = 0\,\mu M^{-1}s^{-1}$, illustrate how the VASP density along the filament changes in each case. **f,** The measured cofilin occupation probability along the filament $P_c$ (y-axis) plotted against the zyxin-VASP self-affinity ($k^v_{\text{bond}}$) (x-axis) for all simulations in the two phase diagrams in (Fig. 3 **c,d**). The dashed black line marks the value of $P_c$ of Eq. 4-5 where $v_{growth} = v_{shrink}$, corresponding to the analytical value which predict the transition between the *depolymerization treadmilling state*. **g,** Cumulative number of polymerization and depolymerization events per filament plotted over time. The slope of each curve is then proportional to the polymerization and depolymerization velocities. **h,** The mean interface deformation $\langle L - L_0\rangle$ as a function of the zyxin-VASP self-affinity $k^v_{\text{bond}}$. System size $15\,\mu m$.

**Extended Data Table 1 Conversion of units between simulation and experimental values.** Summary of the rate conversion of simulation values compared to experimental values. ED stands for Extended Data.

| Parameter | simulation value in numeric units | simulation value in physical units | literature value/ experiments |
|---|---|---|---|
| Lattice length $l_a$ | 1 | 250 $nm$ | - |
| Simulation time $s_{sim}$ | 1 | 10 $s$ | - |
| Actin concentration (G- and F-actin) | 24000 | 1 $\mu M$ | 2 $\mu M$ |
| VASP concentration | 8000 | 0.33 $\mu M$ | 0.125-1 $\mu M$ Fig. 4 |
| cofilin concentration | 24000 | 1 $\mu M$ | 0.5-1 $\mu M$ Fig. 4 |
| Attachment rate VASP on actin $k^{\text{v}}_{\text{on}}$ | 6.5 | 12 $\mu M^{-1}s^{-1}$ | 10 $\mu M^{-1}s^{-1}$ [2, 3] |
| Detachment rate VASP on actin $k^{\text{v}}_{\text{off}}$ | 2 | 1 $s^{-1}$ | 1-4 $s^{-1}$ [2, 3] |
| Attachment rate cofilin on actin $k^{\text{c}}_{\text{on}}$ | 1.2 | 2 $\mu M^{-1}s^{-1}$ | 0.171 $\mu M^{-1}s^{-1}$ [4] |
| Detachment rate cofilin on actin $k^{\text{c}}_{\text{off}}$ | 1.2 | 0.6 $s^{-1}$ | 0.135 $s^{-1}$ [4] - 0.6 $s^{-1}$ [5] |
| Polymerization rate $k_{\text{pol}}$ | $3 \cdot 10^{-4}$ | 72 $\mu M^{-1}s^{-1}$ | 34 $\mu M^{-1}s^{-1}$ ED Figure 5d |
| Depolymerization rate $k_{\text{dis}}$ | 10 | 100 $s^{-1}$ | 53 $s^{-1}$ [6] |
| VASP Bond formation $k^{\text{v}}_{\text{bond}}$ (phase separation) | 0.1 | 8 $\mu M^{-1}s^{-1}$ | - |
| VASP Bond breakage $k_{\text{break}}$ | 2 | 1 $s^{-1}$ | - |
| Persistence length of actin (off-axis growth) | 3.5 | 4 $\mu m$ | 10 $\mu m$ [7] |
| Actin alignment (overlap penalty) | 4.5 | - | - |
| Maximal VASP concentration (steric repulsion) | 5 | 1.3 $\mu M$ | - |

### Supplemental Information

#### Supplementary Note 1: Treadmilling Condition

Among the three states discussed in the main text, filaments are treadmilling in the *treadmilling state* and within a droplet, in the *trapped actin state*, but not in the *depolymerization state*. The key feature of the latter state is that filaments' disassembly proceeds too quickly compared to the filaments' polymerization. This can be formalized by requiring that, for treadmilling to happen, on average, the shrinkage velocity at the back of a filament must be lower than the growth velocity at the tip: $v_{\mathrm{pol}} > v_{\mathrm{dis}}$. As described in Supplementary Note 5, for a polymerization event to occur, a VASP agent must be present at the site, along with a G-actin monomer in the bulk. Consequently, the growth velocity is proportional to the average VASP-density concentration within a bundle $\phi_{\mathrm{v}}$ and the concentration of G-actin monomers $\phi_{\mathrm{a}}^{\mathrm{G}}$, assumed to be homogeneous in the bulk since the monomers are diffusing in 3D. Disassembly, on the other hand, requires the presence of a cofilin molecule at the back of the filament. Therefore, the disassembly velocity is proportional to the cofilin occupation probability $P_{\mathrm{c}}$ along the filament. In terms of the parameters of the system, we can rewrite the two velocities as follows:

$$v_{\mathrm{pol}} = k_{\mathrm{pol}}\, \phi_{\mathrm{v}}\, \phi_{\mathrm{a}}^{\mathrm{G}} \tag{1}$$

$$v_{\mathrm{dis}} = k_{\mathrm{dis}}\, P_{\mathrm{c}} \tag{2}$$

where $k_{\mathrm{pol}}$ and $k_{\mathrm{dis}}$ are tunable rates. If initially $v_{\mathrm{growth}} > v_{\mathrm{shrink}}$ holds, filaments on average grow, increasing their length and reducing the concentration of G-actin monomers $\phi_{\mathrm{a}}^{\mathrm{G}}$, until a balance between disassembly and polymerization velocity is reached $v_{\mathrm{pol}} = v_{\mathrm{dis}}$ (Extended Data Fig. 8g). If, on the other side, the system starts in a regime of $v_{\mathrm{pol}} < v_{\mathrm{dis}}$, the filaments only shrink and no treadmilling is possible. To check whether this treadmilling condition holds for our simulations, we reformulated it as a condition for the cofilin occupation probability

$$P_{\mathrm{c}} < \phi_{\mathrm{v}} \phi_{\mathrm{a}}^{\mathrm{G}} k_{\mathrm{pol}} / k_{\mathrm{dis}} \tag{3}$$

where $P_{\mathrm{c}}$ and $\phi_{\mathrm{v}}$ can be measured directly from the simulations, while $\phi_{\mathrm{a}}^{\mathrm{G}}$ represents the initial value of G-actin concentration. As can be seen in Extended Data Fig. 8f, this threshold accurately separates simulations showing treadmilling from those which do not. It's important to note, that the threshold value for $P_{\mathrm{c}}$ jumps up when zyxin-VASP self-affinity $k_{bond}^{v}$ is high enough to induce phase separation, as $\phi_{\mathrm{v}}$ goes from a value close to the homogeneous density concentration to the binodal value (Extended Data Fig. 8e). This explains why the line characterizing the threshold for treadmilling becomes so sharp (Fig. 3c,d) at very low zyxin-VASP self-affinity values.

The cofilin occupation probability $P_{\mathrm{c}}$ can also be determined as the steady-state solution of the time-dependent distribution $P_{\mathrm{c}}(t)$. To solve for this, we introduce a master equation describing the relevant reactions along the filament. Assuming that only the attachment/detachment dynamics are relevant, $P_{\mathrm{c}}$ is coupled solely to the VASP occupation probability $P_{\mathrm{v}}$. As the model allows for more filaments to overlap on the same lattice site, the master equations contains the occupation probability for each site of each of the $n$ filaments:

$$\frac{d}{dt} P_{\mathrm{v},i} = -k_{\mathrm{off}}^{\mathrm{v}} P_{\mathrm{v},i} + (1 - P_{\mathrm{v},i} - P_{\mathrm{c},i}) k_{\mathrm{on}}^{\mathrm{v}} \phi_{\mathrm{v}}^{\mathrm{free}} \tag{4}$$

$$\frac{d}{dt} P_{\mathrm{c},i} = -k_{\mathrm{off}}^{\mathrm{c}} P_{\mathrm{c},i} + (1 - P_{\mathrm{v},i} - P_{\mathrm{c},i}) k_{\mathrm{on}}^{\mathrm{c}} \phi_{\mathrm{c}}^{\mathrm{free}} \tag{5}$$

where $P_{\alpha,i}$ represents the probability of species $\alpha \in \{\mathrm{c},\mathrm{v}\}$ of filament $i \in \{1, \dots n\}$ and . The first term in both equations represents a loss event, where a protein bound to the filament unbinds, while the second term represents a gain event, with an empty site becoming occupied by a VASP or cofilin agent respectively. $\phi_{\mathrm{v}}^{\mathrm{free}}$ and $\phi_{\mathrm{c}}^{\mathrm{free}}$ are the concentrations of unbound VASP and cofilin, which are free to attach to the filament. As unbound cofilin can diffuse in the bulk, its membrane-density value quickly relaxes to a homogeneous distribution $\phi_{\mathrm{c}}^{\mathrm{free}} = \phi_{\mathrm{c}}$. On the other side, the diffusivity of VASP is limited by its anchoring to the membrane and the overall density $\phi_{\mathrm{v}}$ is a sum of unbounded VASP concentration $\phi_{\mathrm{v}}^{\mathrm{free}}$ and the bound VASP concentration. The bound VASP concentration can be extracted from the set of probabilities $P_{\mathrm{v},j}$. In particular, under the condition that the filaments $i$ is unoccupied, is new $\frac{1}{l_a^2} \sum_{j \neq i} P_{v,j}$ where $l_a$ is the lattice spacing. Rescaling space, such that the lattice spacing is set to unity, we find: $\phi_v^{free} = \phi_v - \sum_{i \neq j} P_{v,i}$, where the sum is over all other filaments on the site. Inserting the value of $\phi_v^{free}$ and $\phi_c^{free}$ in 4–5, we get the final expressions for the master equations for $P_{c,i}$ and $P_{v,i}$. The average concentrations of VASP localized to the bundles was numerically measured to be $\phi_v = 4.2$ particles per lattice, which corresponds to $1.2\,\mu M$, if converted into physical units (Appendix C). This value allows analytically computing the steady-state solution of the cofilin occupation probability of Eqs. (4)-(5), which is found to decrease with increasing $\phi_v$, $k^c_{\text{off}}$ and $k^v_{\text{on}}$ while increasing with higher $k^c_{\text{on}}$ and $k^v_{\text{off}}$, as expected. Comparing the numerically measured $P_c$ with the steady state solutions of Eqs.(4)-(5) for different $n$, we find that the two-filaments approximation well represents the region in parameter space close to the onset of treadmilling, and correctly identifies the threshold where $v_{\text{shrink}} = v_{\text{growth}}$ (Extended Data Fig. 8f).

#### Supplementary Note 2: Transition between the trapped actin and the treadmilling state

To investigate the transition from the *trapped actin* state to a *treadmilling* state, we performed simulations exploring how filament bundles influence VASP condensate interfaces. The simulations presented here are not intended to match specific biochemical parameters of the experimental proteins but rather serve as a theoretical analysis to identify the dominant physical parameters governing condensate–filament interactions. We initialized the system with a bundle of three filaments oriented orthogonally to a straight VASP interface and examined whether the bundle could grow by deforming the interface. Simulation results across parameter ranges (Extended Data Fig. 8a,b) reveal that interface deformation is governed by three main factors: the surface tension of the VASP condensate, filament stiffness, and the distribution of VASP/cofilin along filaments. Surface tension sets the energetic penalty for interface deformation, and is determined by the zyxin-VASP self-affinity $k^v_{\text{bond}}$ (Supplementary Note 4), which describes how fast new bonds between VASP agents are formed. Our results show that sufficiently low tension is required for deformation to occur. Filament stiffness, quantified by the filaments' persistence length, determines whether filaments bend and align with the interface or maintain their orientation. If stiffness is low, filaments tend to align with the interface and grow inside the VASP condensate without deforming it. Conversely, higher stiffness allows filaments to push against and deform the interface. This explains why, in the simulation, single filaments tend to align with the interface, while filament bundles, which have higher stiffness, can deform the zyxin-VASP condensate.

Last, high cofilin occupancy—arising at high cofilin concentrations—prevents VASP attachment (due to their competition) and accelerates filament shrinkage, both of which hinder interface deformation. Consistent with this, we find that increasing cofilin concentration reduces deformation.

In summary, surface tension and cofilin occupancy oppose interface deformation, while filament stiffness promotes it. As shown in Extended Data Fig. 8a—c, simulations in the treadmilling regime produce mean interface deformations exceeding $1.5\mu m$, which sets a numerically inferred threshold for bundle formation in the full system. When filaments deform the zyxin-VASP condensate into a bundle-like shape and these bundles connect into a percolating network, the continuous movement of filaments is enabled, and the system is in the *treadmilling state*. In contrast, in the trapped actin regime, filaments fail to deform the interface and remain confined within the VASP droplet. This transition represents a dynamical analog of the energetic competition between surface tension and filament stiffness discussed in [8].

Our findings contribute to the growing understanding of how condensates and cytoskeletal filaments influence each other's morphology and dynamics. By explicitly modeling the mechanical coupling between filament growth and condensate interface deformation, we address open questions raised in recent reviews [9, 10] in particular, how filament networks reshape condensates and how condensates regulate filament assembly. While our numerical analysis identifies key physical parameters such as surface tension and filament stiffness, it only hints at the underlying mechanisms. A more complete theoretical framework is now needed to understand this dynamic interplay.

#### Supplementary Note 3: Conversion of simulation parameter values into experimental units

At each time step, the probability of a reaction occurring is determined by its rate, as explained in the Methods. Therefore, the intrinsic units of a simulation depend on the specific code's architecture. In particular, the simulation's time unit $s_{sim}$ depends on the fixed time step $dt_{sim}$, lengths are measured in terms of the lattice side $l_a$, and densities $\phi^{sim}$ in the number of agents per area: $[\phi^{sim}] = N^{sim}[n \times l_a]^{-2}$, where $n$ is the number of lattice sites per side of the simulation grid and $N^{sim}$ the number of agents. For a quantitative comparison between simulations and experiments, simulation units have to be converted into physical units and this conversion will be carried trough in this section. As the simulations contain only the minimal set of coarse-grained reactions of the experimental setting, the purpose of the unit's conversion is not to make quantitative comparisons or predictions about the physical system, but rather to ensure that the results and the physical effects observed in our simulations well describe the phenomenology of the experiments. To carry the conversion in physical units, we must choose the level of coarse-graining. We assume that 1 agent in the simulations corresponds to 100 proteins in the experimental setting, fixing the conversion between the number of proteins in the experimental setting $N^{exp}$ and the number of agents in the simulations $N^{sim}$ to

$$N^{exp} = 100 N^{sim}. \tag{6}$$

For simplicity, we consider the same conversion rate for all protein species. This choice can be further used to fix the conversion of space units. Using that an actin monomer has a length of 2.77nm, that reduces to 2.07nm after binding of cofilin [5], we deduce that an actin agent has a length of $200 - 300nm$. As the actin monomers size in the simulations exactly corresponds to the lattice spacing, we obtain the conversion rate for the space units:

$$l_a = 250nm \tag{7}$$

The last unit to fix is the time scale. This is done by requiring that the typical time needed for a VASP protein to detach from the filament match between simulations and experiments, as will be carried out in the next subsection.

#### Conversion of Detachment Rates

In this subsection we show how to fix the timescale of the simulations, done by matching the typical detachment time for a VASP protein between simulations and experiments. A similar conversion metric could be obtained using the detachment rate of cofilin instead, without significant differences. Note that a detachment rate is a convenient choice to fix the time conversion rate, as it only contains a time unit. However, converting the detachment rate of a cofilin or VASP protein from an actin monomer presents some challenges. The difficulties arise because simulations provide a coarse-grained representation of the physical system, where each simulation agent corresponds to 100 real proteins.

In particular, the typical detachment time $\tau_1^{\text{sim}}$ for a single agent in the simulation corresponds, in the physical system, to the detachment time $\tau_{100}^{\text{exp}}$ for 100 proteins. Since individual detachment events occur independently, the detachment time for all 100 proteins corresponds to the slowest detachment time in a statistical ensemble of 100 realizations of the single-agent detachment process:

$$\tau_{100}^{\text{exp}} = \max_{i=1,\ldots,100} (\tau_1^{\text{exp}})_i, \tag{8}$$

where $i$ indexes different stochastic realizations of the same process. By formulating a master equation for $P_n(t)$, which represents the probability that $n$ agents have detached from the filament at time $t$, we derive that:

$$\tau_{100}^{\text{exp}} = (\ln(100) + 1)\tau_1^{\text{exp}}. \tag{9}$$

The master equation for $P_n(t)$ is given by:

$$P_n(t + dt) = P_0(dt)P_n(t) + (100 - n + 1)P_1(dt)P_{n-1}(t) - (100 - n)P_1(dt)P_n(t). \tag{10}$$

Since the probability of multiple detachment events within an infinitesimal time $dt$ scales as $(dt)^m$ for $m \geq 2$, we neglected higher-order terms and only considered single detachment events. The initial condition is set such that at $t = 0$, no detachment event has yet occurred: $P_0(0) = 1$.

Each term on the right-hand side of the equation has a specific meaning: (i) the first term represents the scenario where no detachment event occurs within $[t, t + dt]$, (ii) The second term accounts for a gain in the number of detached proteins, when a system with $n - 1$ detached agents transitions to $n$, (iii) The third term accounts for a loss, when the system transitions from $n$ to $n - 1$. The probability for a single detachment event or for no event in an infinitesimally small time $dt$ can be given in terms of linear rates:

$$P_1(dt) = k_{\text{off}}^{\text{exp}} dt. \tag{11}$$

Here, $k_{\text{off}}^{\text{exp}}$ is the inverse of the typical detachment time. Neglecting higher order terms the probability of no detachment events occurring is then given by:

$$P_0(dt) = 1 - k_{\text{off}}^{\text{exp}} dt$$

Inserting these expressions into the master equation and expanding to first order in $dt$, we obtain the differential equation:

$$\frac{dP_n(t)}{dt} = -k_{\text{off}}^{\text{exp}}(100-n)P_n(t) + k_{\text{off}}^{\text{exp}}(100-n+1)P_{n-1}(t). \tag{12}$$

To derive the typical time $\langle t \rangle_n$ required to observe $n$ unbindings, we integrate over time:

$$-1 = k_{\text{off}}^{\text{exp}}(101-n)\left(\langle t\rangle_{n-1} - \langle t\rangle_n\right). \tag{13}$$

Solving this recursion equation gives:

$$\langle t\rangle_n = \frac{1}{k_{\text{off}}^{\text{exp}}}\sum_{i=1}^{n}\frac{1}{101-i}. \tag{14}$$

Setting $n = 100$, we obtain the total detachment time for 100 proteins:

$$\tau_{100}^{\text{exp}} = \frac{1}{k_{\text{off}}^{\text{exp}}}\sum_{i=1}^{100}\frac{1}{i} \approx \frac{1+\ln(100)}{k_{\text{off}}^{\text{exp}}}. \tag{15}$$

Since we require that the typical detachment time of one agent in simulation matches 100 detachment events in physical units ($\tau_1^{\text{sim}} = \tau_{100}^{\text{exp}}$), we obtain the conversion rate for the detachment rate:

$$k_{\text{off}}^{\text{exp}} = (\ln(100)+1)k_{\text{off}}^{\text{sim}}. \tag{16}$$

Here, the left-hand side is in seconds ($s^{-1}$), while the right-hand side is in simulation time units ($s_{\text{sim}}^{-1}$). To determine the conversion rate for time, we require that the literature value for the detachment rate of cofilin ($k_{\text{off/cofilin}}^{\text{exp}} = 0.6s^{-1}$, [5]) matches our conversion metric. This gives:

$$0.6\frac{1}{s} = 1.2(\ln(100)+1)\frac{1}{s_{\text{sim}}}. \tag{17}$$

Solving for $s_{\text{sim}}$, we obtain:

$$s_{\text{sim}} = 11.2s \approx 10s. \tag{18}$$

where we have used the detachment rate of VASP in the simulation ($k_{\text{off/cofilin}}^{\text{sim}} = 1.2s_{\text{sim}}^{-1}$).

#### Concentrations

In this section, we show how to convert the concentration values of VASP, cofilin and actin from numerical to physical units. The conversion of concentration values is needed to convert attachment rates of VASP or cofilin agents to the filament, as will be carried out in the next subsection. The main conceptual difficulty when converting concentrations lies in the fact that the simulations are performed on a 2-dimensional lattice, while the experiments are intrinsically 3-dimensional. To address this issue, we follow a similar strategy as done in [11] and introduce an additional length scale $l_b$, which represents the height of the solution above the lipid bilayers. The additional length scale $l_b$ is a free parameter in our conversion. We set this length equal to the average distance traveled by a protein in solution in the simulation time $s_{sim} = 6s$ (see next section):

$$l_b = \sqrt{D\,dt_{sim}} = 18\mu m \approx 10\mu m \tag{19}$$

where the diffusion coefficient of a protein in the solution was estimated using the Stokes-Einstein relation $D = k_bT/(6\pi\eta R) \approx 8\mu m^2 s^{-1}$, where $T$ was taken to be the room temperature $298\,K$, the radius of the protein $R$ for cofilin or G-actin is around $3\,nm$, the viscosity around $1\,mPa$, similar as in the experiments. The length scale $l_b$ enables the conversion of simulation concentration values, here expressed in physical units, into experimental concentration values:

$$\phi_{exp} = \frac{N_{exp}}{(l_a)^2 l_b} = \frac{100N_{sim}}{(l_a)^2 l_b} = \frac{100}{l_b}\phi_{sim} \tag{20}$$

where $\phi_{exp/sim}$ are the concentration values respectively for experiments and simulations, $N_{exp/sim}$ the number of molecules/agents per site and the factor 100 comes from Eq. 6. The concentrations in physical units for the simulations are summarized in Table 1, with densities expressed in micromoles using the relation $1\mu M = 600\text{particle}/\mu m^3$.

#### Attachment rate to an actin filament

The theoretical attachment rates of VASP and cofilin to the filament can be derived similarly to their detachment rates. In our simulations, which provide a coarse-grained representation of the experimental system, a single attachment event of an agent corresponds to the attachment of 100 proteins in reality. The time required for all 100 proteins to attach is determined by the slowest attachment event in an ensemble of 100 independent realizations of a single-agent attachment. As a result, the scaling factor for attachment follows the same form as that for detachment, given by $\ln(100) + 1$ (see Eq. 16).

A key distinction between attachment and detachment processes is that the attachment rate is concentration-dependent. Consequently, the conversion must also incorporate the concentration scaling. The relationship for converting attachment rates from simulation units to physical units is given by:

$$k_{\text{on}}^{\text{exp}} = \frac{l_b}{100} k_{\text{on}}^{\text{sim}} (\ln(100) + 1), \tag{21}$$

where the factor $100/l_b$ accounts for the concentration conversion (see Eq. 20). The converted attachment rates are summarized in Table 1.

The cofilin attachment rate used in the simulations, when converted to physical units, is approximately two orders of magnitude higher than the experimentally measured value for single cofilin binding. However, this is consistent with the coarse-grained nature of the model, which represents cooperative binding of multiple cofilin molecules rather than single-molecule kinetics. Experimental cooperativity coefficients for cofilin binding range from 2.3 to 20. Additionally, CAP1 enhances cofilin's depolymerization efficiency and competes for filament binding by interacting with F-actin sides [6]. To incorporate these effects, we model CAP1's influence as an effective increase in the cofilin attachment rate.

#### Persistence length of actin filaments

In the numerical model, filaments' stiffness is incorporated by favoring the probability $P_s$ of straight polymerization, over the probability $P_t$ of polymerization in one of the two tilted directions. Assuming that a growth event occurs, such that $P_s + P_t = 1$, the probability of polymerization in a tilted direction is penalized over straight growth, but can occur in two different directions, leading $P_t = 2e^{-\mu_{bend}} P_s$.

This preference for straight growth leads to the filament persistence length $l_p$ of the filament, defined as the length over which the filament maintains its direction before undergoing a change in orientation. To infer $l_p$ from the off-axis growth penalty $\mu_{bend}$, we calculate the average length after which a bending event happen:

$$l_p = l_a \sum nP_n = l_a \frac{P_s P_t}{(1 - P_s)^2} = l_a \frac{e^{\mu_{bend}}}{2} \tag{22}$$

where $P_n = (P_s)^n P_t$ is the probability that the filament has grown straight $n$ times before changing its direction. Converting the above expression from the length simulation unit $l_a$ to $\mu m$, gives the value reported in Table 1. In simulations, the persistence length is approximately four times smaller than that observed in experiments without cofilin [7]. However, it is known that the binding of cofilin to actin filaments reduces filament stiffness and thus the persistence length to a value comparable to that in our simulations [12]. Note also that the persistence length of filaments in a bundle cross-linked by VASP scales linearly with the number of filaments in the bundle.

#### Further remarks

The polymerization and depolymerization rates, denoted as $k_{growth}$ and $k_{shrink}$, can also be converted to physical units, similarly as for the attachment rate ( Eq. (21)). However, instead of using the scaling factor $\ln(100) + 1$, the factor is simply 100. This is because polymerization and depolymerization of 100 molecules occur sequentially, not independently. In addition, to properly convert the polymerization rate $k_{\text{growth}}$, an additional rescaling with the system size is required. The reason is that, in the simulations, the effective rate for a growth event is the polymerization rate $k_{pol}$ multiplied by the total number of free monomers in the bulk. Hence, it scales extensively with the system size. This differs from the experimental situation, where a growth event scales with the local monomer concentration and is therefore intensive with respect to the system size. However, in our simulations the bulk concentration of G-actin is homogeneous, such that the conversion between local density and the total amount of G-actin can be readily inferred. Rewriting the effective probability rate of a growth event gives: $k_{\text{grow}}^{\text{sim}} N_{\text{actin}}^{\text{sim}} = k_{\text{grow}}^{\text{sim}} L_{\text{sim}}^2 \phi_{\text{actin}}^{\text{sim}}$, where $L_{\text{sim}}$ is the system size. Absorbing the factor $L_{\text{sim}}^2$ into the definition of $k_{\text{grow}}^{\text{sim}}$ yields an intensive parameter that corresponds to the experimental value. Therefore, we obtain the following conversions for the rates:

$$k_{shrink}^{\text{exp}} = 100\, k_{shrink}^{\text{sim}} \quad \text{and} \quad k_{grow}^{\text{exp}} = L_{sim}^2 l_b\, k_{grow}^{\text{sim}}$$

$$k_{grow}^{\text{exp}} = L_{sim}^2 l_b\, k_{grow}^{\text{sim}} = (80)^2 (10\mu m)(3 \cdot 10^{-4} \frac{(l_a)^2}{N_{g-actin} s_{sim}})$$

$$k_{grow}^{\text{exp}} = L_{sim}^2 l_b\, k_{grow}^{\text{sim}} = (80)^2 (10\mu m)(3 \cdot 10^{-4} \frac{(l_a)^2}{N s_{sim}}) = (80)^2 \frac{10}{10 \cdot 16} (3 \cdot 10^{-4}) 600\ \mu M^{-1} s^{-1} = 72 \mu M^{-1} s^{-1}$$

On the other hand, the rates of bond formation and breakage between VASP agents are more difficult to coarse-grain. The number of bonds between agents changes during the coarse-graining process, and is not constant. Furthermore, the experimental values for these bond formation and breakage rates are unknown, making a direct comparison impossible. For this reason, we do not attempt the coarse-graining procedure for these rates. Instead, we simply express them in physical units to provide an understanding of the scales involved at the coarse-grained level.

Following the same approach as for the attachment and detachment rates, we have the following conversions:

$$k_{break}^{\text{exp}} = k_{break}^{\text{sim}} \quad \text{and} \quad (k_{\text{bond}}^{v})^{\text{exp}} = \frac{16 l_b}{100} (k_{\text{bond}}^{v})^{\text{sim}}$$

The factor of 16 in the $k_{\text{bond}}^{v}$ conversion accounts for the fact that the interaction between two VASP agents scales with the number of bonds they can form.

#### Supplementary Note 4: Surface tension of the VASP condensate

In our simulations, VASP phase separation is modeled through bond formation between neighboring VASP agents. Specifically, two adjacent VASP agents can form a bond with rate $k_{bond}^{v}$, while an existing bond can break with rate $k_{break}$. Once formed, a bond cannot be stretched by diffusion of either agent; consequently, the mobility of bonded VASP agents is effectively reduced. When many bonds are formed, a VASP condensate emerges. The aim of this section is to demonstrate that the parameter $k_{bond}^{v}$ controls the formation of the VASP condensate and governs the surface tension of condensate's interface.

Although our model is formulated in terms of stochastic reaction rates rather than an explicit free-energy functional, the reversible VASP–VASP bonding dynamics are thermodynamically consistent at equilibrium. In particular, bond formation and bond breaking obey detailed balance. Denoting by $p_{\text{bound}}$ and $p_{\text{unbound}}$ the probabilities that two neighboring VASP agents are in the bound and unbound state, respectively, detailed balance requires

$$k_{bond}^{v}\, p_{\text{unbound}} = k_{break}\, p_{\text{bound}}. \tag{23}$$

It follows that

$$\frac{p_{\text{bound}}}{p_{\text{unbound}}} = \frac{k_{bond}^{v}}{k_{break}} \equiv K. \tag{24}$$

At thermodynamic equilibrium, the ratio of probabilities of two microscopic states is given by a Boltzmann weight,

$$\frac{p_{\text{bound}}}{p_{\text{unbound}}} = \exp\left(-\frac{\Delta G_{\text{contact}}}{k_B T}\right), \tag{25}$$

where $\Delta G_{\text{contact}}$ denotes the free-energy difference between a VASP–VASP contact and the corresponding VASP–background contact. Combining the above relations yields

$$K = \frac{k_{bond}^{v}}{k_{break}} = \exp\left(-\frac{\Delta G_{\text{contact}}}{k_B T}\right). \tag{26}$$

Therefore, the kinetic ratio $K$ directly encodes the effective interaction free energy between VASP molecules. Increasing $K$ corresponds to making VASP–VASP contacts thermodynamically more favorable relative to

VASP–background contacts. In mean-field descriptions of phase separation, such as Flory–Huggins theory, demixing is controlled by an effective interaction parameter $\chi$ that quantifies the relative favorability of protein–protein versus protein–background contacts. The relation above shows that $K$ plays an analogous role in our kinetic model: increasing $K$ increases the effective attraction strength and thus enhances the thermodynamic driving force toward phase separation. Since $k_{break}$ is kept fixed throughout our simulations, varying $k_{bond}$ directly varies $K$. For this reason, we use $k^v_{bond}$ (or equivalently $K$) as an operational measure of what we refer to as the *VASP self-affinity*. To further show numerically that $k^v_{bond}$ controls condensation, we performed simulations in the absence of actin and cofilin, with VASP initially confined to the left half of the system (Supplementary Movie 5). This setup creates an initial interface between a high-density condensate region and a low-density region. We observe that for $k^v_{bond} < k_c$, the interface gradually dissolves and the system evolves toward a homogeneous VASP density. In contrast, for $k^v_{bond} > k_c$, the interface remains stable over time. These results are consistent with Fig. 3c,d, where VASP condensation is observed at similar parameter values. Taken together, these observations indicate that $k^v_{bond}$ controls VASP condensation and effectively plays a role analogous to a Flory-Huggins interaction parameter $\chi$.

Next, we quantify how $k_{bond}$ affects the surface tension $\gamma$ of the condensate once it has formed. To do so, we measure the difference between the instantaneous interface length $L$ and the length of the corresponding straight interface $L_0$. This difference is non-zero because thermal fluctuations deform the initially straight interface, generating additional interfacial length. The creation of excess length is opposed by the surface tension $\gamma$, which tends to minimize the total interface length.

We will show that the time-averaged excess interface length is governed by the surface tension according to

$$\langle L - L_0 \rangle_t \propto \frac{k_B T}{\gamma}. \tag{27}$$

The measured values of $\langle L - L_0 \rangle_t$ are shown in Extended Data Fig. 8h and the corresponding simulations are shown in Supplementary Movie 5. We find that increasing $k^v_{bond}$ leads to a monotonic reduction in the mean interface fluctuations. Using Eq. 27, this implies that the surface tension $\gamma$ increases monotonically with $k_{bond}$.

To justify Eq. 27, we briefly outline its derivation within capillary wave theory. As the interactions between VASP proteins are purely passive, we assume that the system is ergodic, such that the time average $\langle \cdot \rangle_t$ is equivalent to the ensemble average $\langle \cdot \rangle$. We parameterize the deviations from a straight interface by a height function $h(y,t)$ in the Monge gauge, which assumes no overhangs in the interface profile. The initial straight interface corresponds to $h(y,0) = 0$. Within this parametrization, ensemble averages can be expressed as path integrals over $h(y)$ weighted by the Boltzmann factor $e^{-\beta \mathcal{F}[h]}$, where $\beta = (k_B T)^{-1}$.

The excess interface length can be written as

$$\langle L - L_0 \rangle = \left\langle \int_0^{L_0} dy \left[ \sqrt{1 + (\partial_y h)^2} \right] - L_0 \right\rangle. \tag{28}$$

Assuming weak perturbations from a straight line, $(\partial_y h)^2 \ll 1$, we expand to leading order:

$$\langle L - L_0 \rangle \approx \left\langle \int_0^{L_0} \frac{(\partial_y h)^2}{2} \, dy \right\rangle. \tag{29}$$

We now express the height field in Fourier space$h(y) = \sum_q h_q e^{iqy}$, which gives:

$$\langle L - L_0 \rangle \approx \left\langle \int_0^{L_0} \frac{(\partial_y h)^2}{2} \, dy \right\rangle = \sum_q q^2 \langle |h_q|^2 \rangle. \tag{30}$$

To evaluate this expression, we specify the interfacial free energy using the standard capillary wave form:

$$\mathcal{F}[h] = \frac{\gamma}{2} \int dy \, (\partial_y h)^2 = \frac{\gamma}{2} \sum_q q^2 |h_q|^2. \tag{31}$$

Since the free energy is quadratic in the Fourier modes, each independent mode contributes $\frac{1}{2}k_BT$ on average by the equipartition theorem. Therefore,

$$\frac{\gamma}{2}q^2\langle|h_q|^2\rangle = \frac{1}{2}k_BT, \tag{32}$$

which implies

$$q^2\langle|h_q|^2\rangle = \frac{k_BT}{\gamma}. \tag{33}$$

Substituting this result into Eq. 30, we obtain

$$\langle L - L_0\rangle \propto \frac{k_BT}{\gamma}, \tag{34}$$

which establishes Eq. 27.

### Supplementary Note 5: Detailed Code Description

All agents are updated sequentially according to a fixed index list chosen at the start of the simulation and kept constant during the simulation. We consider periodic boundary conditions. Initially, we randomly place $N_{\text{vasp}}$ VASP agents, $N_{\text{cofilin}}$ cofilin agents, and $N_{\text{filament}}$ filaments with an initial length of $L_0 = 7$ and random orientation. This mimicks experimental conditions where the initial lack of aged ADP-actin prevents cofilin binding, allowing fast early polymerization. The number of these three entities is conserved throughout the simulation. From the initial state, we evolve the system through diffusion and reaction events. Each biochemical process $i$ is associated with a rate $k_i$, so that the probability for the process to occur, within a small interval $dt$, is $k_idt$. The corresponding waiting time $\tau_i$ is exponentially distributed as $p(\tau_i) = k_i \exp(-k_i\tau_i)$. The process with the smallest waiting time is executed, and the simulation clock advances by $dt$. In the simulations used throughout this study, $dt = 0.001$. However, we tried to lower $dt$ till $dt = 0.00001$ and checked that we obtained consistent results.

#### VASP

For every VASP agent that is not filament-bound, diffusion is always sampled. A waiting time $\tau_D$ is drawn for hopping to each neighboring site containing fewer than 5 VASP agents; steric exclusion is enforced only for VASP, since they are the largest components. In addition, a neighboring site is considered eligible only if moving the VASP agent there does not stretch any of its existing bonds with other VASP agents, i.e. all bonded partners must remain nearest neighbors after the hop. This effectively reduces the mobility of clustered agents relative to individual ones, as observed experimentally. If at least one eligible site yields $\tau_D < dt$, the agent diffuses to the site associated with the smallest $\tau_D$. Independently of diffusion, an unbound VASP agent can undergo exactly one of the following mutually exclusive reaction events within a given time step: filament attachment, bond formation, or bond dissociation. A waiting time $\tau_i$ is sampled for each admissible event type; if none satisfies $\tau_i < dt$, no reaction occurs. Otherwise, the reaction with the smallest $\tau_i$ is executed. A bond can form only with VASP agents on the same or neighboring sites, provided both agents have fewer than 4 occupied bonds. For each potential partner, a waiting time $\tau_{\text{bond}}$ is sampled from an exponential distribution with rate proportional to the product of the two agents' numbers of free bonds, implemented as $k^v_{\text{bond}}n_{\text{free bonds}}$. Bond breakage occurs with a rate $k^{\text{v}}_{\text{break}}n_{\text{bonds}}$, proportional to the number of existing bonds $n_{\text{bonds}}$. Attachment is possible only if free F-actin monomers (not occupied by cofilin or VASP) are present at the agent's site. The attachment waiting time $\tau_{\text{attach}}$ is sampled from an exponential distribution with rate $k^{\text{v}}_{\text{on}}n_{\text{free sites}}$, where $n_{\text{free sites}}$ is the number unbounded actin monomers on the same lattice site of the VASP agent.

For a filament-bound VASP agent, 2D diffusion is not allowed. Instead, within each $dt$ the agent can perform at most one of the following mutually exclusive events: detach, hop forward, or hop backward. Forward hopping is forbidden at the filament tip, while backward hopping is forbidden at both the tip and the back of the filament. The capture mechanism at the filament tip mimicks the preferential adhesion of VASP to barbed ends observed experimentally [13]. Waiting times for all admissible events are sampled from their respective exponential distributions, and if any satisfies $\tau_i < dt$ , the event with the smallest $\tau_i$ is executed; otherwise no event occurs.

#### Cofilin

For a cofilin agent that is not filament-bound, only attachment to a filament is explicitly sampled. Possible attachment sites are restricted to sites that (i) coincide with the cofilin position, and (ii) are empty (no other particles are attached). For each eligible filament site, a reaction time for attachment $\tau_{\mathrm{attach}}$ is sampled from the exponential distribution associated with the attachment rate $k^{\mathrm{c}}_{\mathrm{on}}$. If the minimum reaction time satisfies $\tau_{\mathrm{attach}} < dt$, the cofilin agent attaches to the filament site corresponding to the smallest sampled reaction time. If no attachment event occurs within $dt$, the cofilin agent is repositioned to a random site, which represents fast three-dimensional diffusion projected onto the two-dimensional simulation plane. Note that the cooperative binding of cofilin is only taken into account in the explicit value of $k^{\mathrm{c}}_{\mathrm{on}}$, but is not implemented as a reaction, as our system represents a mesoscopic description of the experiments (1 agent corresponds to 100 particles in the experiment).

For a filament-bound cofilin agent, detachment is the only possible process. A reaction time $\tau_{\mathrm{detach}}$ is sampled from the exponential distribution associated with the detachment rate $k^{\mathrm{c}}_{\mathrm{off}}$. If $\tau_{\mathrm{detach}} < dt$, the agent detaches from the filament and is subsequently repositioned to a random site, again modeling fast 3D diffusion. If $\tau_{\mathrm{detach}} \geq dt$, the agent remains bound and no event is executed during that time step.

#### Filament

If a filament has cofilin bound at its minus end and its length $L$ is bigger than two, $L > 2$, a shrinkage event can occur. Filaments with length $L \leq 2$ are considered inactive ("dead"): they do not undergo shrinkage but can still elongate by polymerization. Growth of a dead filament beyond this length threshold effectively re-activates it; we refer to this transition as nucleation. This convention allows us to treat the system as approximately filament-number conserving while still permitting filament death and nucleation events. Accordingly, filaments shorter than three monomers are not shown in the videos or included in the analysis. When a disassembly event is allowed, a reaction time $\tau_{\mathrm{dis}}$ is sampled from the exponential distribution associated with the disassembly rate $k_{\mathrm{dis}}$. If $\tau_{\mathrm{dis}} < dt$, the filament shortens by one monomer at the minus end, and the pool of free actin monomers is increased by one.

Depolymerization (disassembly) is treated independently from filament growth, i.e. a filament may shrink and still be evaluated for growth within the same update. Filament growth is evaluated at the plus end by considering three possible growth sites, corresponding to the allowed growth directions.

Filament polymerization is VASP-catalyzed only, i.e. spontaneous polymerization is not included. This mimics experimental conditions in which profilin and capping protein suppress spontaneous filament growth, so that elongation occurs predominantly through VASP-mediated incorporation. For each candidate growth site, eligible VASP agents are chosen. A VASP is eligible to catalyze growth only if it sits exactly on the candidate growth site and is not filament-bound. One single VASP agent is sufficient to catalyze growth. Three possible sites for growth are possible, corresponding to the three forward directions from the head where the filament sits. For each eligible VASP–site pair, a reaction time $\tau_{\mathrm{pol}}$ is sampled from the exponential distribution associated with a growth rate proportional to $k_{\mathrm{pol}}$ and to the number of free actin monomers on the site $n_{\mathrm{free\ actin}}$, and modulated by geometric penalty factors. In particular, straight growth is preferred via an off-axis growth penalty, reducing the probability rate to grow in one of the two bent directions. Explicitly, the rate of off-axis growth is $k_{\mathrm{pol}} n_{\mathrm{free\ actin}} e^{-\mu_{bend}}$. An additional overlap penalty is applied when the candidate growth configuration would overlap with another existing filament. In this case, the growth rate is reduced by a factor of $e^{-\mu_{cross}}$. Among all candidate growth events (across the three possible growth sites and all eligible VASP agents), the event with the shortest sampled reaction time is selected. If this reaction time satisfies $\tau_{\mathrm{pol}} < dt$, the filament grows by one monomer at the selected plus-end site. The selected VASP agent is then attached to the newly created filament site, and the pool of free actin monomers is decreased by one. Note that by increasing the number of free monomers during shrinkage events and decreasing it during growth events, we ensure that the total number of actin monomers—whether in the G-actin or F-actin state—is conserved throughout the simulation. Taken together, an actin filament can move only through polymerization and depolymerization, and thermal fluctuations are not taken into account, as zyxin-mediated anchoring to the membrane and the formation of cross-linked bundled filament structures suppress fluctuations (Supplementary Movie 1).

We do not explicitly model ATP-to-ADP aging within F-actin. Since cofilin induces depolymerization only at the pointed end in the experimental system (no fragmentation), nucleotide state would affect turnover only through the terminal subunit. Under the experimental conditions considered, the ATP/ADP-Pi cap is expected to span only a small fraction of the filament length, while simulated filaments are typically much longer. Therefore, the terminal subunit is effectively in the ADP state on the turnover timescale, and neglecting explicit nucleotide aging does not qualitatively affect filament dynamics or regime classification. This was confirmed using exemplary simulations including a subunit ageing step.

#### Supplementary Movie Captions

**Supplementary Movie 1. Actin polymerization by zyxin-VASP condensates.** Zyxin-VASP condensates (blue) ($C_{zyxin-VASP} = 0.25\,\mu M$) polymerize actin (red hot) ($C_{actin} = 2\,\mu M$) from condensate clusters as seen in Fig. 1d. This forms a network of aligned filaments for zyxinΔLIM-VASP and disordered filaments for zyxinFL-VASP. After stalling of initial polymerization, zyxin-VASP clusters slowly reform. After the initial fast polymerization (0-5 min), the frame rate is increased to account for the slow reformation of zyxin-VASP droplets. Scale bars, 10 µm.

**Supplementary Movie 2. Emergence of treadmilling-like actin bundle movement by competing zyxinΔLIM-VASP and cofilin.** Competition of cofilin and CAP1 ($C_{cofilin} = 0.5\,\mu M$, $C_{CAP1} = 0.125\,\mu M$) with the zyxinΔLIM-VASP decoration ($C_{zyxin\Delta LIM-VASP} = 0.5\,\mu M$) on actin bundles (red hot) leads to directional movement of bundles. Bundles are simultaneously polymerized and oppositely disassembled. The zyxinΔLIM-VASP condensates remains colocalized to the bundles during the whole process. At bundle crossover events, zyxinΔLIM-VASP-condensates are transiently depleted from the bundles by the barbed ends of polymerizing bundles. Scale bars, 10 µm.

**Supplementary Movie 3. Actin network states in simulations.** Simulations varying the self-affinity strength of zyxin-VASP result in the treadmilling state ($k^v_{\text{bond}} = 8\,\mu M^{-1}s^{-1}$), the depolymerization state ($k^v_{\text{bond}} = 3\,\mu M^{-1}s^{-1}$), and the trapped actin state ($k^v_{\text{bond}} = 17\,\mu M^{-1}s^{-1}$). The square domain is $20\,\mu m$ per side. Actin filaments are shown in orange and the density of zyxin-VASP agents in blue, while the density of cofilin is not shown. The other parameter values used in this simulation are summarized in Extended Data Table 1.

**Supplementary Movie 4. Coarsening of bundles of filaments in simulations.** Simulation showing the coarsening of filament bundles over time, square domain of $60\,\mu m$ per side. The zyxin-VASP complex phase-separates so that the simulation is within the treadmilling state (parameter regime same as in Supplementary Movie 3 with $k^v_{\text{bond}} = 8\,\mu M^{-1}s^{-1}$).

**Supplementary Movie 5. Zyxin–VASP self-affinity modulates the surface tension of the condensate.** The simulation shows that the mean interface deformation of the zyxin–VASP interface (solid black line) decreases with increasing zyxin–VASP self-affinity $k^v_{\text{bond}}$. System size, $15\,\mu m$.

**Supplementary Movie 6. Enhanced treadmilling-like movement velocity by high cofilin concentrations.** Increasing cofilin concentration from $C_{cofilin} = 0.5\,\mu M$ to $C_{cofilin} = 1\,\mu M$, competing with the zyxin-VASP condensates ($C_{zyxin\Delta LIM-VASP} = 0.25\,\mu M$) (blue), dramatically increases velocity of treadmilling-like movement and bundle coarsening dynamics of the actin filaments (red hot). Scale bars, 10 µm.

**Supplementary Movie 7. High zyxinΔLIM-VASP concentrations lead to jammed actin dynamics.** At $C_{zyxin\Delta LIM-VASP} = 1\,\mu M$, the high density of filaments on the bilayer arrest bundle movement in a jammed state. Treadmilling-like bundle movement emerges only in short burst from $+1/2$ nematic defects. These dynamics are enhanced at higher cofilin concentrations ($C_{cofilin} = 1\,\mu M$) with a reemergence of bundle coarsening. Scale bars, 10 µm.

**Supplementary Movie 8. Treadmilling-like dynamics are limited by surface tension of zyxinFL-VASP condensates.** Additional interactions in zyxinFL-VASP condensates (blue) decouple the actin dynamics (red hot) from the condensates at a concentration of $C_{zyxinFL-VASP} = 0.5\,\mu M$ ($C_{cofilin} = 0.5\,\mu M$, $C_{CAP1} = 0.125\,\mu M$) (Fig. 5a–c). Solely at precisely $C_{zyxinFL-VASP} = 0.075\,\mu M$, directional treadmilling-like movement of small actin bundles is possible (Fig. 5f). Scale bars, 10 µm.
